# An unsuspected physiological role for mGluRIII glutamate receptors in hippocampal area CA1

**DOI:** 10.1101/2025.03.31.646479

**Authors:** Maurice A. Petroccione, Marcello Melone, Travis J. Rathwell, Namit Dwivedi, Christine Grienberger, Fiorenzo Conti, Annalisa Scimemi

**Affiliations:** SUNY Albany, Department of Biology, 1400 Washington Avenue, Albany (NY) 12222 – USA; Università Politecnica delle Marche, Department of Experimental and Clinical Medicine, Via Tronto 10/a, Ancona 60126 – Italy; IRCCS INRCA, Center for Neurobiology of Aging, Via Birarelli 8, Ancona 60121 – Italy; IISER Pune, Dr. Homi Bhabha Road, Pune 411008 – India; Brandeis University, Department of Biology and Volen National Center for Complex Systems, Waltham (MA) 02453 – USA

**Keywords:** Metabotropic glutamate receptors, mGluRIII, hippocampus, presynaptic, inhibition, place cells, CA1 spatial navigation, *in vivo* patch clamp

## Abstract

Group III metabotropic glutamate receptors (mGluRIII) are expressed broadly throughout the neocortex and hippocampus but are thought to inhibit neurotransmitter release only at a subset of synapses and in a target cell-specific manner. Accordingly, previous slice physiology experiments in hippocampal area CA1 showed that mGluRIII receptors inhibit glutamate and GABA release only at excitatory and inhibitory synapses formed onto GABAergic interneurons, not onto pyramidal cells. Here, we show that the supposed target cell-specific modulation of GABA release only occurs when the extracellular calcium concentration in the recording solution is higher than its physiological concentration in the cerebrospinal fluid. Under more physiological conditions, mGluRIII receptors inhibit GABA release at synapses formed onto both interneurons and pyramidal cells but limit glutamate release only onto interneurons. This previously unrecognized form of mGluRIII-dependent, pre-synaptic modulation of inhibition onto pyramidal cells is accounted for by a reduction in the size of the readily releasable pool, mediated by protein kinase A and its vesicle-associated target proteins, synapsins. Using *in vivo* whole-cell recordings in behaving mice, we demonstrate that blocking mGluRIII activation in the intact CA1 network results in net effects consistent with decreased inhibition and significantly alters CA1 place cell activity. Together, these findings challenge our current understanding of the role of mGluRIII receptors in the control of synaptic transmission and encoding of spatial information in the hippocampus.

## Introduction

Group III metabotropic glutamate receptors (mGluRIII) are a family of G-protein coupled receptors expressed widely throughout the brain. Ultrastructural studies in rats indicate that mGluRIII receptors are mainly localized in pre- synaptic terminals and axonal compartments^1–13^. Their primary role is to depress the release of glutamate, GABA, and neuromodulators, such as acetylcholine, monoamines, and peptides^1–13^. In the hippocampus, two types of mGluRIII receptors with high affinity for glutamate, mGluR4a (EC_50_=3-20 µM) and mGluR8a (EC_50_=0.02 µM), are expressed pre-synaptically, at both excitatory (E) and inhibitory (I) synaptic contacts^9,10,12,14,15^. By contrast, mGluR7a, a low-affinity mGluRIII type (EC_50_=1 mM), is thought to be expressed more abundantly at E compared to I pre-synaptic terminals and at pyramidal cell-to-interneuron (PC-IN) more than at pyramidal cell-to-pyramidal cell (PC-PC) synapses^11^. Consistent with this ultrastructural evidence, previous electrophysiology experiments in acute hippocampal slices showed that E/I synapses onto INs are more sensitive to mGluRIII modulation than E/I synapses formed onto PCs^3,16^. Because these effects could be detected in the presence of mGluRIII agonists like L-AP4 and in conditions that promote glutamate spillover (e.g., glutamate uptake blockers, repetitive stimulations), it was suggested that the mGluRIII-mediated inhibition of neurotransmitter release is activity-dependent and occurs only when the extracellular concentration of glutamate is increased from its baseline level^3,16^. Since mGluRIII receptors are coupled predominantly to G_i/o_ proteins, their pre-synaptic I-effect was thought to be mediated by inhibition of adenylyl cyclase, but more detailed information on the molecular mechanisms underlying this modulatory effect remained unclear.

When reviewing past slice physiology literature, an important methodological consideration is that many experiments have been performed using so-called “standard” extracellular recording solutions^17^, where the extracellular calcium concentration ([𝐶𝑎^2+^]_𝑜_) was set to 2-2.5 mM. This concentration range, which is more than 2- fold higher than what we now know to be present in the cerebrospinal fluid under physiological conditions (∼1.2 mM)^18^, was often chosen to facilitate neurotransmitter release, allowing the recording of larger currents^19^. Whether this masked an mGluRIII-dependent modulation of neurotransmitter release at E/I synapses onto CA1-PCs, to the best of our knowledge, has not been previously explored.

Here, we show that mGluRIII receptors are expressed at a subset of E/I synapses onto CA1-PCs and that under experimental conditions that mimic more closely the extracellular calcium concentration of the cerebrospinal fluid, the target cell-specificity of the mGluRIII modulation of GABA release onto CA1-PCs is no longer detected. mGluRIII activation reduces synaptic inhibition from parvalbumin- and somatostatin-positive INs (PV- and SST-INs, respectively). By contrast, the target-cell specific modulation of excitation by mGluRIII activation persists in [𝐶𝑎^2+^]_𝑜_=1.2 mM, leading to a reduced excitation of CA1-INs but not CA1-PCs. Finally, we examined the network effects of mGluRIII activation *in vivo,* showing that it limits location-specific firing of CA1-PCs, with important implications for their ability to generate accurate representations of spatial information in the hippocampus.

## Results

### mGluRIII receptors are expressed at E/I synapses onto CA1-PCs

To analyze the sub-cellular distribution of mGluRIII in the mouse hippocampus, we used a pre-embedding electron microscopy (EM) approach and identified the ultrastructural localization of mGluR4a, mGluR7a, and mGluR8a at E- and I-synapses onto CA1-PCs, in the PC layer and *stratum radiatum*. The immunoreactivity for these receptors was localized in axons, axon terminals forming both asymmetric (glutamatergic, E) and symmetric (GABAergic, I) synaptic contacts, dendrites and astrocytic processes (**Figure 1A-B; Table S1**). All mGluRIII receptor types were expressed pre- and post-synaptically (**Figure 1A-B**). We compared the degree of expression of each mGluRIII type at asymmetric versus symmetric synapses, to determine whether their abundance varies between E/I synapses onto CA1-PCs (**Figure 1C**). We then measured the proportion of mGluRIII immuno-positive and immuno-negative axon terminals onto CA1-PCs (mGluRIII+ and mGluRIII-, respectively; **Figure 1C**, **Table 1**) and found that, out of these three mGluRIII isoforms, mGluR7a was the only one that displayed a preferential pre-synaptic expression at I-synapses (***p=7.0e-4). By contrast, mGluR4a and mGluR8a showed a comparable degree of pre-synaptic expression at E- and I-synapses onto CA1-PCs (p=0.07, and p=0.09, respectively; **Table 1**, **Figure 1C**).

**Figure 1.**
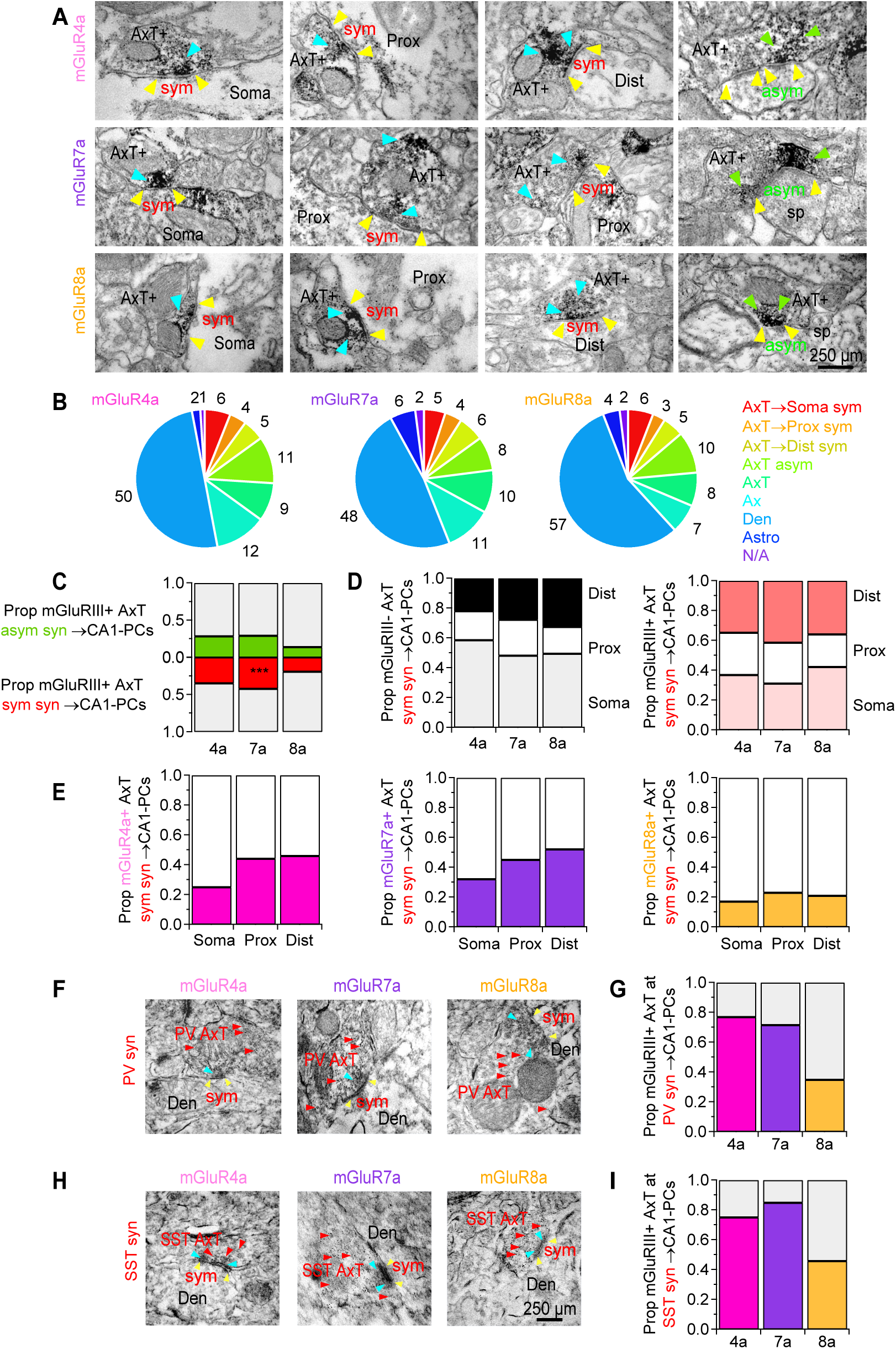
Sub-cellular distribution of mGluRIII receptors at E/I synapses onto CA1-PCs. **(A)** Electron micrographs showing immuno-reactivity for mGluR4a, mGluR7a and mGluR8a (*cyan arrowheads*) in the axon terminal (*AxT*) of asymmetric and symmetric synapses (*yellow arrowheads*) targeting the soma, proximal (*Prox*) and distal dendrites (*Dist*), and dendritic spines (*sp*) of CA1-PCs. **(B)** Pie charts showing the quantification of the immuno-labeling EM data. All mGluRIII receptors were expressed post- and pre-synaptically, especially at I-synapses onto CA1-PCs. **(C)** Summary of the proportion of E/I synapses onto CA1-PCs that are immuno-positive for different mGluRIII types. mGluR7a is more abundant at I than E synapses onto CA1-PCs. **(D)** *Left,* Summary of the spatial distribution of I-synapses immuno-negative for different mGluRIII types. *Right,* Summary of the spatial distribution of I-synapses immuno-positive for different mGluRIII types. **(E)** Proportion of analyzed I-synapses that are immuno-positive and -negative for mGluR4a (*left*), mGluR7a (*center*) and mGluR8a (*right*). **(F)** Electron micrographs showing immuno-reactivity for mGluR4a, mGluR7a and mGluR8a (*cyan arrowheads*) in the axon terminal (*AxT*) of symmetric synapses formed by PV-INs (*red arrowheads*) onto the dendrites (*Den*) of CA1-PCs. **(G)** Proportion of immuno-positive PV synapses out of all those that were analyzed. **(H-I)** As in F-G for synapses formed by SST-INs onto CA1-PCs.

**Table 1.** Comparison of mGluR4a, mGluR7a, and mGluR8a localization at symmetric and asymmetric synapses onto CA1-PCs.

| Target | AxT profiles | AxT+<br>sym | AxT-<br>sym | AxT+<br>asym | AxT-<br>asym | sym vs asym |
| --- | --- | --- | --- | --- | --- | --- |
| *mGluR4a* | n=743 | 35%<br>n=141 | 65%<br>n=263 | 29%<br>n=97 | 71%<br>n=242 | p=0.07 |
| *mGluR7a* | n=699 | 42%<br>n=154 | 58%<br>n=212 | 29%<br>n=89 | 71%<br>n=214 | ***p=7.0e-4 |
| *mGluR8a* | n=662 | 19%<br>n=64 | 81%<br>n=267 | 14%<br>n=47 | 86%<br>n=284 | p=0.09 |

The spatial distribution of mGluR4a+ and mGluR7a+ I-terminals differed from that of mGluR4a- and mGluR7a- I- terminals (***p=1.6e-4 and **p=2.9e-3, respectively; **Table 2**, **Figure 1D**). Most mGluR4a- and mGluR7a- I- terminals were perisomatic (**Figure 1D**, *left*), whereas mGluR4a+ and mGluR7a+ I-terminals were distributed similarly across the perisomatic compartment, proximal and distal dendrites (**Figure 1D**, *right*). mGluR8a+ and mGluR8a- I-terminals were more likely to occur in the soma and distal dendrites compared to proximal dendrites (p=0.55; **Table 2**, **Figure 1D**). Overall, the proportion of mGluR4a+ axon terminals increased from 25% at the soma to 46% in distal dendrites (**Figure 1E**, *left*), and that of mGluR7a+ terminals increased from 32% at the soma to 52% in distal dendrites; **Figure 1E**, *center*). By contrast, the distribution of mGluR8a+ terminals did not vary significantly, being 17% at the soma and 21% in distal dendrites (**Figure 1E**, *right*).

**Table 2.** Comparison of mGluR4a, mGluR7a, and mGluR8a localization at immuno-positive (+) and -negative (-) axon terminals at symmetric synapses targeting the soma, proximal and distal dendrites of CA1-PCs. The percentage values represent the proportion of all immune-positive/negative symmetric synapses present at the soma, proximal and distal dendrites. The percentage values in parentheses represent the proportion of all somatic, proximal, and distal symmetric synapses immuno-positive/negative for each mGluRIII isoform.

| Target | AxT profiles | sym+/- | AxT>Soma | AxT>Prox | AxT>Dist | Difference across compartments |
| --- | --- | --- | --- | --- | --- | --- |
| *mGluR4a* | n=404 | sym+<br>n=141<br>sym-<br>n=263 | 37%<br>n=52<br>(25%)<br>59%<br>n=154<br>(75%) | 28%<br>n=40<br>(44%)<br>19%<br>n=51<br>(56%) | 35%<br>n=49<br>(46%)<br>22%<br>n=58<br>(54%) | ***p=1.6e-4 |
| *mGluR7a* | n=366 | sym+<br>n=154<br>sym-<br>n=212 | 31%<br>n=48<br>(32%)<br>48%<br>n=102<br>(68%) | 27%<br>n=42<br>(45%)<br>24%<br>n=51<br>(55%) | 42%<br>n=64<br>(52%)<br>28%<br>n=59<br>(48%) | **p=2.9e-3 |
| *mGluR8a* | n=331 | sym+<br>n=64<br>sym-<br>n=267 | 42%<br>n=27<br>(17%)<br>49%<br>n=132<br>(83%) | 22%<br>n=14<br>(23%)<br>18%<br>n=48<br>(77%) | 36%<br>n=23<br>(21%)<br>33%<br>n=87<br>(79%) | p=0.55 |

Two major types of GABAergic interneurons (INs) in the hippocampus, the parvalbumin (PV) and somatostatin (SST) INs, preferentially innervate the soma/proximal dendrites and distal dendrites of CA1-PCs, respectively ^20^. By using a double immuno-peroxidase and pre-embedding immunogold labeling approach in tissue sections from *Pvalb*^Cre/+^:*Gt(ROSA)26Sor*^tm9/+^ and *Sst*^Cre/+^:*Gt(ROSA)26Sor*^tm9/+^ mice, we found that most I-terminals from PV- and SST-INs onto CA1-PCs express mGluR4a/7a (PV-INs: 71-77%; SST-INs: 75-85%), whereas only a subset of them expresses mGluR8a (PV-INs: 35%; SST-INs: 46%; **Figure 1F-I**).

Together, these findings indicate that: *(i)* all mGluRIII receptor types are expressed pre- and post-synaptically; *(ii)* mGluRIII receptors are expressed pre-synaptically at a subset of E/I pre-synaptic terminals onto CA1-PCs; *(iii)* mGluRIII+ I-terminals target the soma, proximal and distal dendrites of CA1-PCs; *(iv)* mGluR4a and mGluR7a are the most abundant mGluRIII isoforms expressed at I-synapses onto CA1-PCs; *(v)* most I-synapses onto CA1-PCs formed by PV- and SST-INs are mGluR4a+ and mGluR7a+, with a smaller proportion expressing mGluR8a.

### mGluRIII activation reduces inhibition onto CA1-PCs at physiological ***[𝑪𝒂^𝟐+^]_𝒐_***

Despite ultrastructural evidence of mGluRIII expression at subsets of E/I synapses onto CA1-PCs, these receptors are thought to inhibit glutamate and GABA release only at E/I synapses targeting CA1-INs, not CA1-PCs, based on experimental evidence collected under conditions that increase the concentration of glutamate in the extracellular space ^3,16^. This conclusion, however, was reached based on experiments performed using solutions containing [𝐶𝑎^2+^]_𝑜_ =2.5 mM. We confirmed this finding in our experiments, in which: *(i)* we used [𝐶𝑎^2+^]_𝑜_ =2.5 mM; (*ii*) we increased the extracellular glutamate concentration by blocking GLT-1, the most abundant glutamate transporter in the adult brain, with DHK (100 µM) ^21^; and (*iii*) we evoked IPSCs in CA1-PCs by delivering electrical stimuli to *stratum radiatum*. As shown in **Figure 2A-C**, DHK did not lead to significant changes in the IPSC amplitude and kinetics when the [𝐶𝑎^2+^]_𝑜_=2.5 mM, in agreement with previous work ^16^ (amp: 93±14%, p=0.66; t_50_: 105±10%, p=0.62; rise: 96±10%, p=0.73, n=10). Since this [𝐶𝑎^2+^]_𝑜_ is more than 2-fold higher than the physiological [𝐶𝑎^2+^]_𝑜_ in the cerebrospinal fluid (i.e., 1.2 mM) ^18^, we repeated the experiments in [𝐶𝑎^2+^]_𝑜_=1.2 mM (**Figure 2D-F**). In this case, DHK reduced the IPSC amplitude in CA1-PCs to 69±6% of its baseline value (n=15, ***p=8.8e-5) without inducing any significant change in the IPSC decay and rise time (t_50_: 93±4%, p=0.13; rise: 100±7%, p=0.96). To confirm that mGluRIII receptors mediated this effect, we applied DHK in the continued presence of the mGluRIII antagonist MSOP (100 µM). Under these conditions, DHK did not change the IPSC amplitude, decay, or rise time (amp: 98±9%, p=0.82; t_50_: 91±5%, p=0.15; rise: 103±16%, p=0.88, n=7; **Figure 2G-I**). These findings suggest that: *(i)* blocking glutamate transporters promotes mGluRIII activation and *(ii)* in contrast to what previously thought, when [𝐶𝑎^2+^]_𝑜_ is within the physiological range, spillover-mediated mGluRIII activation reduces GABAergic inhibition onto CA1-PCs. To determine whether this effect could be attributed to a particular type of mGluRIII receptor with very low affinity for glutamate and abundantly expressed in the CA1 area, like mGluR7a, we applied DHK in the continued presence of the mGluR7a antagonist MMPIP (10 µM). Under these conditions, DHK did not change the IPSC amplitude, decay or rise time (amp: 101±7%, p=0.85; t_50_: 100±7%, p=0.94; rise: 112±13%, p=0.35, n=10; **Figure 2J-L**), suggesting that the mGluRIII-dependent pre-synaptic inhibition of GABA release onto CA1-PCs is largely mediated by mGluR7a. For reasons that remain unclear, in slices, mGluRIII activation requires increased extracellular glutamate levels, because blocking mGluRIII receptors with MSOP in the absence of DHK did not change the amplitude and kinetics of IPSCs recorded from CA1-PCs (amp: 92±6%, p=0.23; t_50_: 96±3%, p=0.28; rise: 94±5%, p=0.25, n=10; **Figure 2M-O**). We hypothesize that this might be due to the slicing procedure, which, by severing cortical and intrahippocampal excitatory connections, might lead to a lower extracellular glutamate concentration than the one detected *in vivo*. We addressed this concern later in this work through a series of experiments in awake-behaving mice.

**Figure 2.**
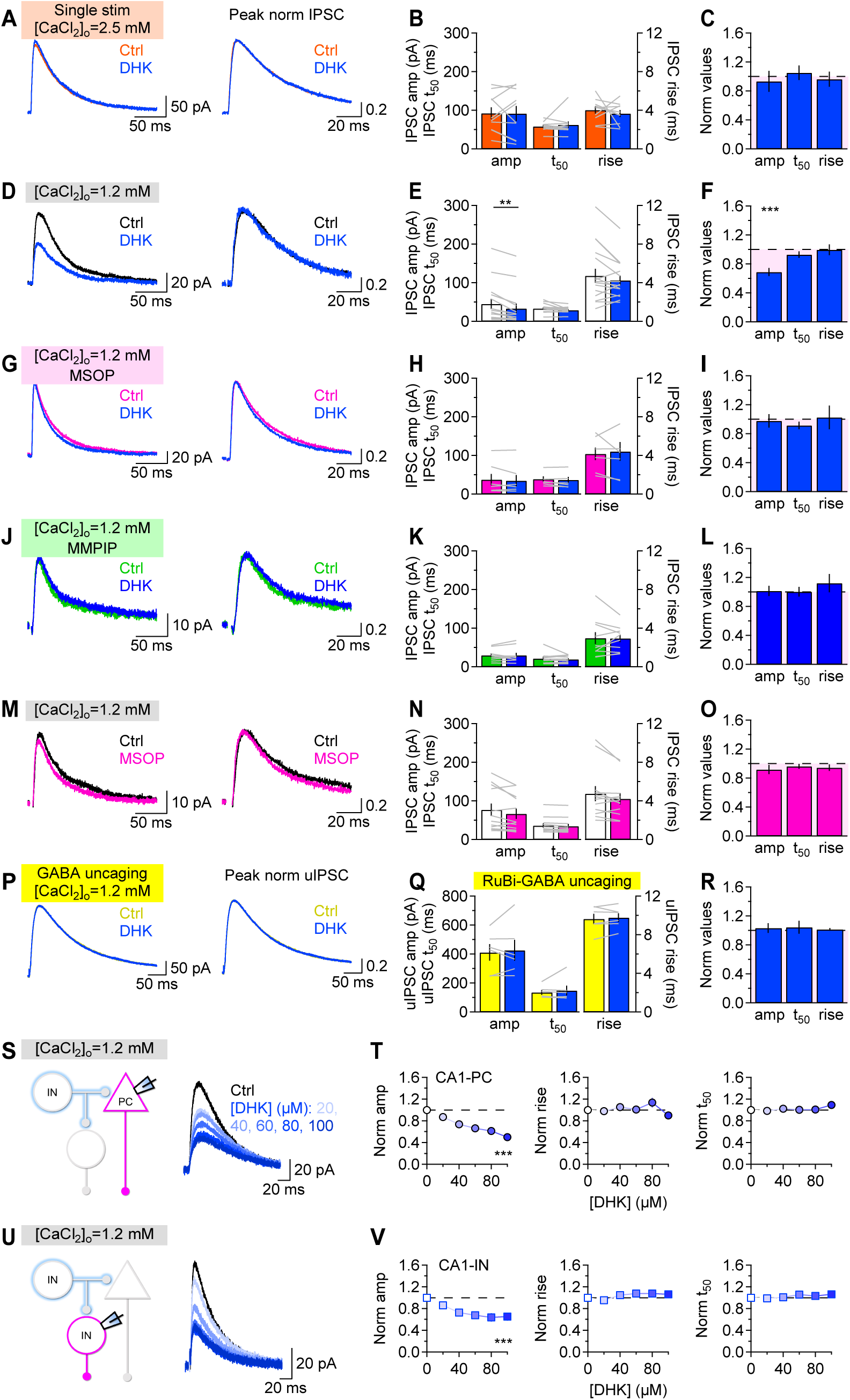
mGluRIII activation reduces inhibition onto CA1-PCs. **(A)** Average IPSCs recorded with [𝐶𝑎]^2+^=2.5 mM in CA1-PCs, before (*orange*) and after DHK (100 µM; *blue*). The traces on the right are normalized by the peak IPSC amplitude. **(B)** Summary of the effect of DHK on the IPSC amplitude and kinetics. **(C)** Summary of the relative change in IPSC amplitude and kinetics induced by DHK. **(D-L)** As in A-C, DHK was applied in the presence of [𝐶𝑎]^2+^=1.2 mM (D-F), and in the continued presence of MSOP (100 µM; G-I) or MMPIP (10 µM; J-L). **(M-O)** As in A-C, in baseline conditions and in the presence of MSOP. **(P-R)** As in A-C, for uIPSCs evoked by uncaging RuBi-GABA (5 µM) in [𝐶𝑎^2+^]_𝑜_=1.2 mM and in the presence of TTX (1 µM), NBQX (10 µM) and APV (50 µM), before and after DHK. **(S)** Schematic representation of the local circuit activated by the electrical stimulation when recording IPSCs from CA1-PCs. The traces represent average IPSCs recorded in the presence of increasing concentrations of DHK. **(T)** Summary of the effect of different DHK concentrations on the amplitude, rise and decay of IPSCs recorded from CA1-PCs. **(U-V)** As in S-T, for IPSCs recorded from CA1-INs in *stratum radiatum*.

To determine whether our findings held true throughout the neocortex, we repeated our experiments in principal cells of the primary somatosensory cortex (SS1-PCs). An mGluRIII-dependent suppression of heterosynaptic inhibition was still detected, suggesting that this modulation of inhibition is not limited to the CA1 region but also occurs in other cortical domains (**Figure S1**).

To rule out the possibility that the effect of DHK might be confounded by activation of post- rather than pre- synaptic mGluRIII, we performed a series of RuBi-GABA uncaging experiments (n=6; **Figure 2P-R**). In these experiments, blocking GLT-1 promotes the activation of both pre- and post-synaptic mGluRIII. However, since GABA is not released from pre-synaptic terminals, a reduction in the amplitude of uncaging evoked IPSCs (uIPSCs) induced by DHK would point to a post-synaptic contribution of mGluRIII, whereas the lack of effect of DHK on the uIPSC amplitude would suggest that activation of postsynaptic mGluRIII does not account for the results shown in **Figure 2D-F**. To test this hypothesis, we blocked action potential (AP) propagation with TTX (1 µM) and evoked uIPSCs by uncaging RuBi-GABA (5 µM) onto CA1-PCs in the presence of ionotropic glutamate receptor blockers (NBQX 10 µM and APV 50 µM). Under these experimental conditions, DHK did not change the uIPSC amplitude and kinetics (amp: 103±7%, p=0.64; t_50_: 105±9%, p=0.63; rise: 102±2%, p=0.42, n=6; **Figure 2M-O**). This suggests that the heterosynaptic depression of GABAergic inhibition onto CA1-PCs, induced by DHK and prevented by the simultaneous application of DHK and MSOP, is mediated by pre-synaptic mGluRIII.

We then asked whether GABAergic inhibition onto CA1-PCs and INs might have a different sensitivity to glutamate uptake blockade in [𝐶𝑎^2+^]_𝑜_=1.2 mM. To answer this question, we monitored the effect of increasing concentrations of DHK (20-100 µM) on IPSCs recorded from CA1-PCs (n=23; **Figure 2S-T**) and *stratum radiatum* INs (n=19; **Figure 2U-V**). The results showed that DHK reduced the IPSC amplitude similarly in CA1-PCs and INs (p=0.98) without changing the IPSC kinetics (CA1-PC: rise p=0.97, t_50_ p=0.99; CA1-INs: rise p=0.23, t_50_ p=0.49). Together, these findings suggest that mGluRIII receptors do not inhibit GABA release in a target cell-specific manner when the [𝐶𝑎^2+^]_𝑜_=1.2 mM, but reduce GABAergic inhibition similarly, onto both CA1-PCs and CA1-INs.

The expression of mGluRIII can also be detected at E-synapses onto CA1-PCs, and even in this case, a target-cell specific effect of mGluRIII activation has been detected in [𝐶𝑎^2+^]_𝑜_=2.5 mM ^3,22^. For this reason, we asked whether DHK could induce a target cell-specific change in the EPSC amplitude in [𝐶𝑎^2+^]_𝑜_=1.2 mM. However, in [𝐶𝑎^2+^]_𝑜_=1.2 mM, DHK did not alter the EPSC amplitude, decay and rise time in CA1-PCs (amp: 100±16%, p=0.98; t_50_: 108±4%, p=0.07; rise: 107±9%, p=0.46, n=10; **Figure S2A-D**), but induced a significant reduction of the EPSC amplitude in CA1-INs (amp: 42±5%, ***p=1.4e-5; t_50_: 100±16%, p=1.00; rise: 98±16%, p=0.90, n=8; **Figure S2E- H**). Similar results were obtained in [𝐶𝑎^2+^]_𝑜_=2.5 mM: DHK did not alter the EPSC amplitude, decay and rise time in CA1-PCs (amp: 85±17%, p=0.41; t_50_: 99±5%, p=0.81; rise: 103±3%, p=0.43, n=14; **Figure S2I-L**), but induced a significant reduction of the EPSC amplitude in CA1-INs (amp: 57±13%, ***p=7.4e-3; t_50_: 106±15%, p=0.69; rise: 100±13%, p=0.13, n=10; **Figure S2M-P**). These findings suggest that rising levels of glutamate in the extracellular space can lead to a target cell-specific reduction of excitation in CA1-IN but not CA1-PCs, regardless of [𝐶𝑎^2+^]_𝑜_.

### mGluRIII activation reduces the size of the readily releasable pool

A reduction in the IPSC amplitude in the presence of DHK may be due to changes in the size of the readily releasable pool for GABA (*RRP,* comprised of *N* releases sites), the release probability (*Pr*), or the quantal size (*q*). To gain insights into whether promoting glutamate spillover with DHK led to changes in the quantal parameter *q*, we analyzed its effect on the amplitude of miniature IPSCs (mIPSCs), recorded from CA1-PCs in the presence of TTX (1 µM; **Figure 3A-C**). The results showed that DHK does not change the mIPSC amplitude (97±4%, n=10, p=0.48) or kinetics (t_50_: 100±3%, n=10, p=0.87; rise: 98±6%, n=10, p=0.74; **Figure 3B-C**), suggesting that DHK does not change the quantal size, *q*.

**Figure 3.**
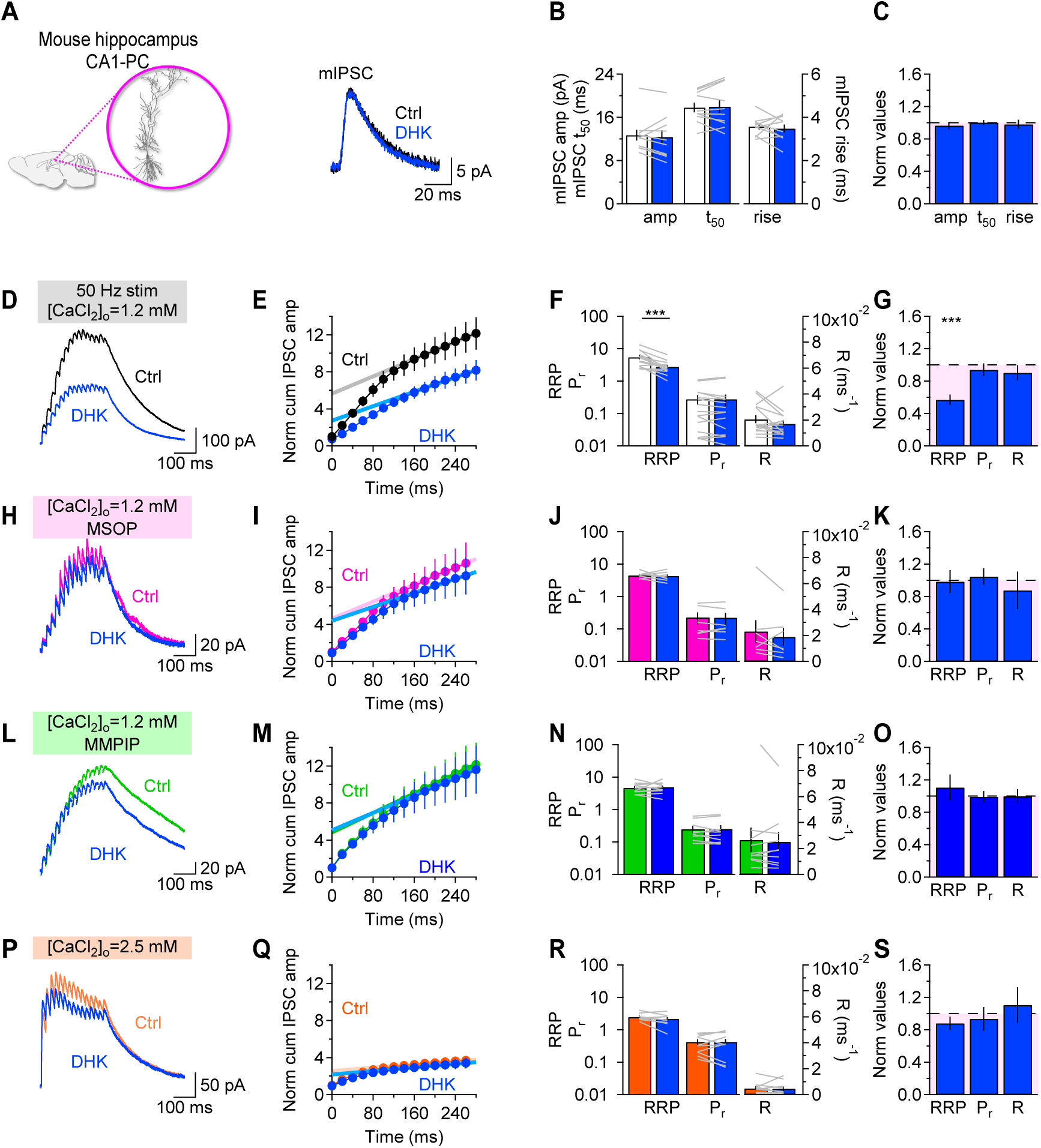
mGluRIII activation reduces the size of the RRP at I-synapses onto CA1-PCs. **(A)** *Left,* Schematic representation of CA1-PCs in sagittal brain slices. *Right,* Average mIPSCs recorded in a CA1-PC in control conditions (*black*) and in DHK (100 µM; *blue*). **(B)** Mean and in-cell comparison of the effect of DHK on the mIPSC amplitude and kinetics. **(C)** Summary of the effect of DHK on the mIPSC amplitude and kinetics, normalized by their values in control conditions. **(D)** Representative IPSC train recorded in response to *stratum radiatum* stimulation with 15 pulses at 50 Hz (280 ms), before (*black*) and after DHK (*blue*), in [𝐶𝑎^2+^]_𝑜_=1.2 mM. **(E)** Cumulative IPSC amplitude normalized by the amplitude of the first IPSC in the train, plotted as a function of the stimulation timing, in control conditions (*black*) and in DHK (*blue*). Thich lines represent the linear fit of the last 5 IPSCs in each condition. The y-intercept of these lines was used to estimate the RRP. **(F)** Mean and in-cell comparison of the effect of DHK on RRP, *Pr* and R. **(G)** Summary of the effect of DHK on RRP, *Pr* and R, normalized by their values in control conditions. **(H-K)** As in D-G, for recordings obtained in the continued presence of the mGluRIII antagonist MSOP (100 µM). **(L-O)** As in D-G, for recordings obtained in the continued presence of the mGluR7a antagonist MMPIP (10 µM). **(Q-S)** As in D-G, for recordings obtained in [𝐶𝑎^2+^]_𝑜_=2.5 mM.

We then delivered a train of 15 pulses at 50 Hz to *stratum radiatum* (**Figure 3D-S**) and estimated the size of the *RRP* using a linear regression of the cumulative IPSC amplitude over time intervals that are short compared to the time required for recovery from synaptic depression ^23–27^. The linear fit of the cumulative IPSC amplitude was performed on the last five IPSCs in the train. The replenishment rate *R* was estimated by the slope of the fit, whereas *Pr* was calculated from the ratio of the IPSC amplitude and the RRP size. The analysis indicated that, when [𝐶𝑎^2+^]_𝑜_=1.2 mM, DHK reduced the *RRP* size without altering *Pr* or the vesicle replenishment/refilling rate ^28–32^(*RRP*_DHK/Ctrl_: 57±6%, n=15, ***p=1.0e-5; *Pr*_DHK/Ctrl_: 94±8%, p=0.45; *R*_DHK/Ctrl_: 91±9%, p=0.33; **Figure 3D-G**). Similar results were also obtained when analyzing IPSCs recorded from SS1-PCs, indicating that similar mechanisms account for heterosynaptic depression of inhibition when glutamate uptake is impaired in different brain areas (**Figure S3**).

The regression analysis does not assume that *Pr* is constant throughout the train or uniform across all vesicles, but a potential caveat is that it assumes constant vesicle replenishment ^33^. To account for potential inconsistencies in vesicle replenishment during the train stimulation, we verified the effect of DHK on the *RRP* using two other methods that rely on different assumptions ^27,34^. For example, the EQ method ^27^ assumes that vesicle replenishment is negligible early during the train, when the synaptic currents decay linearly ^27^. In this case, the *RRP* can be estimated by the *x*-axis intercept of a linear fit in a plot of the IPSC amplitude versus the cumulative IPSC amplitude during the train (**Figure S4A-B**). With the EQ method, the *RRP* in DHK was ∼79% of that in control conditions, with no significant change in the release probability *Pr* (*RRP*_DHK/Ctrl_: 79±5%, n=15, ***p=1.9e-5; *Pr*_DHK/Ctrl_: 89±6%, p=0.08; **Figure S4A-B**).

A third method, referred to as the decay method, also allows estimating the *RRP* size, in this case from an exponential fit of the IPSC amplitude plotted against the stimulus number ^34^. Using the decay method, we still found that DHK reduced the *RRP* size without altering *Pr* (*RRP*_DHK/Ctrl_: 73±11%, n=15, *p=0.4; *Pr*_DHK/Ctrl_: 104±23%, p=0.14; **Figure S4C-D**). Together with the mIPSC analysis, these results indicate that experimental conditions that lead to an increase in the extracellular glutamate concentration induce a reduction of the *RRP* size without changing *Pr* at I-synapses onto CA1-PCs.

To confirm that mGluRIII mediated the effects of DHK, we repeated the train stimulation and the integration analysis on IPSCs recorded in the continued presence of MSOP (100 µM), before and after DHK. Under these conditions, DHK did not change the *RRP* size, release probability and refilling rate (*RRP*_DHK/Ctrl_: 99±14%, n=7, p=0.92; *Pr*_DHK/Ctrl_: 150±10%, p=0.64; *R*_DHK/Ctrl_: 88±23%, p=0.61; **Figure 3H-K**). No change in the *RRP* size, release probability *Pr* and refilling rate *R* were detected when DHK was applied in the continued presence of the mGluR7a antagonist MMPIP, suggesting that the effect of DHK is mostly mediated by mGluR7a receptors (*RRP*_DHK/Ctrl_: 101±11%, n=10, p=0.89; *Pr*_DHK/Ctrl_: 101±7%, p=0.85; *R*_DHK/Ctrl_: 94±8%, p=0.47; **Figure 3L-O**). If mGluRIII did not change inhibition onto CA1-PCs in [𝐶𝑎^2+^]_𝑜_=2.5 mM, we would not expect DHK not to induce any effect on *RRP*, *Pr,* or *R* in [𝐶𝑎^2+^]_𝑜_=2.5 mM. Consistent with this hypothesis, no effect of DHK was detected when [𝐶𝑎^2+^]_𝑜_ was increased from 1.2 mM to 2.5 mM (*RRP*_DHK/Ctrl_: 88±8%, n=10, p=0.17; *Pr*_DHK/Ctrl_: 94±15%, p=0.68; *R*_DHK/Ctrl_: 111±22%, p=0.64; **Figure 3P-S**). Together, these findings suggest that mGluRIII activation limits inhibition onto CA1-PCs by reducing the *RRP* size at GABAergic axon terminals, an effect that only occurs when [𝐶𝑎^2+^]_𝑜_ is in the physiological range.

### mGluRIII receptors reduce the RRP size by reducing PKA and synapsins activation

mGluRIII receptors are G protein-coupled receptors linked to G_i/o_ proteins. In addition to activating K^+^ channels and inhibiting Ca^2+^ channels via liberation of G_βγ_ subunits, they inhibit the membrane-associated enzyme adenylyl cyclase (AC)^13^. By doing so, they reduce the intracellular concentration of cAMP and prevent the activation of protein kinase A (PKA). PKA, in turn, inhibits AC via a negative feedback loop (**Figure S5**)^35,36^. At I-synapses in the cerebellum, PKA inhibition can reduce the size of the *RRP* without altering the release probability *Pr* by changing the phosphorylation state of a family of vesicle-associated proteins called synapsins^37–46^. If PKA-dependent phosphorylation of synapsins is also critical to control the *RRP* size at I-synapses onto CA1-PCs, then the mGluRIII- dependent suppression of inhibition should no longer be detected in the presence of a PKA inhibitor. To test this hypothesis, we recorded IPSCs in the presence of the PKA inhibitor KT5720 (1 µM) before and after DHK (**Figure 4A-G**). KT5720 is an ATP-competitive antagonist for the ATP binding site on the catalytic subunit of PKA ^47,48^. This catalytic subunit must bind ATP before it can phosphorylate Ser/Thr residues on target proteins, so its blockade prevents the cAMP-dependent phosphorylation of PKA substrates ^49^. Under these conditions, DHK did not change the IPSC amplitude or kinetics (amplitude: 95±10%, n=11, p=0.65; t_50_: 99±3%, p=0.85; rise: 109±14%, p=0.54; **Figure 4A-C**), and did not induce any significant change in *RRP* size, *Pr* or the rate of vesicle replenishment of the RRP (*RRP*: 91±6%, n=11, p=0.20; *Pr*: 100±10%, p=0.99; *R*: 105±14%, p=0.76; **Figure 4D-G**). Similar results were also obtained when the IPSCs were recorded from SS1-PCs (**Figure S6**).

**Figure 4.**
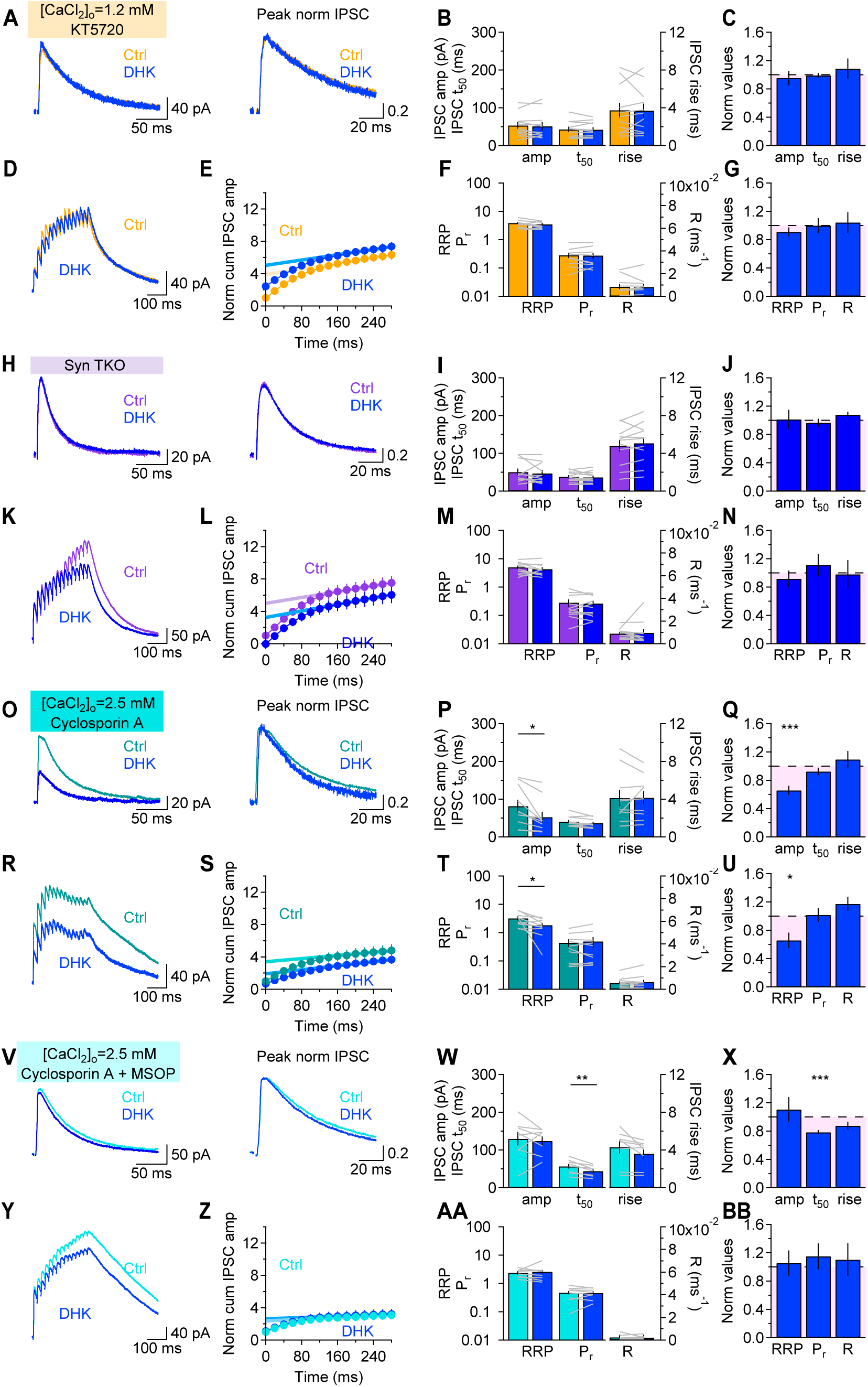
PKA inhibition and synapsins loss prevent the spillover-induced reduction in the RRP at I- synapses onto CA1-PCs. **(A)** Average IPSCs recorded with [𝐶𝑎^2+^]_𝑜_=1.2 mM in CA1-PCs, before (*orange*) and after DHK (100 µM; *blue*), in the continued presence of the PKA inhibitor KT5720 (1 µM). The traces on the right are normalized by the peak IPSC amplitude. **(B)** Summary of the effect of DHK on the IPSC amplitude and kinetics in KT5720. **(C)** Summary of the relative change in IPSC amplitude and kinetics induced by DHK applied in the continued presence of KT5720. **(D)** Representative IPSC train recorded in response to *stratum radiatum* stimulation with 15 pulses at 50 Hz (280 ms), before (*orange*) and after DHK (*blue*), in the continued presence of KT5720 and in [𝐶𝑎^2+^]_𝑜_=1.2 mM. **(E)** Cumulative IPSC amplitude normalized by the amplitude of the first IPSC in the train, plotted as a function of the stimulation timing, in control conditions (*orange*) and in DHK (*blue*). Thich lines represent the linear fit of the last 5 IPSCs in each condition. The y-intercept of these lines was used to estimate the RRP. **(F)** Mean and in-cell comparison of the effect of DHK on RRP, Pr and R, when applied in the presence of KT5720. **(G)** Summary of the effect of DHK on RRP, Pr and R, when applied in the continued presence of KT5720. The data were normalized by their values in control conditions. **(H-N)** As in A-G, for data collected from CA1-PCs in Syn TKO mice, in control conditions (*purple*) and in the presence of DHK (*blue*). **(O-U)** As in A-G, for data collected with [𝐶𝑎^2+^]_𝑜_=2.5 mM and in the continued presence of the calcineurin inhibitor cyclosporin A (1 µM), in control conditions (*teal*) and in the presence of DHK (*blue*). **(V-BB)** As in A-G, for data collected with [𝐶𝑎^2+^]_𝑜_=2.5 mM and in the continued presence of the calcineurin inhibitor cyclosporin A (1 µM) and the mGluRIII antagonist MSOP (100 µM), in control conditions (*cyan*) and in the presence of DHK (*blue*).

If the effects of DHK are mediated by PKA acting on synapsins, then DHK should also not change the IPSC amplitude and *RRP* size in slices from mice in which the genes encoding for synapsins I-III are deleted (here referred to as SynTKO; **Figure 4H-N**). Consistent with this hypothesis, DHK did not change the IPSC amplitude or kinetics in CA1-PCs from SynTKO mice (amplitude: 102±13%, n=11, p=0.91; t_50_: 96±5%, p=0.53; rise: 108±4%, p=0.07; **Figure 4H-J**). In SynTKO mice, DHK did not induce any significant change in *RRP* size, *Pr* or *R* of I-synapses (*RRP*: 92±11%, n=11, p=0.50; *Pr*: 111±16%, p=0.47; *R*: 98±20%, p=0.93; **Figure 4K-N**).

These findings indicate that mGluRIII activation at I-synapses onto CA1-PCs reduces *RRP* through signaling cascades involving reduced PKA activation and reduced vesicle mobilization by synapsins. Given that the modulation of *RRP* at I-synapses onto CA1-PCs is no longer detected at [𝐶𝑎]^2+^=2.5 mM, and *RRP* is smaller in 2.5 compared to 1.2 mM Ca^2+^ (**p=2.1e-3; cf. **Figure 3D-G** and **Figure 3P-S**), we wondered whether any signaling pathway that can change the net phosphorylation state of synapsins in a Ca^2+^-dependent manner may be responsible for the lack of mGluRIII modulation of inhibition in [𝐶𝑎^2+^]_𝑜_=2.5 mM. Multiple protein kinases and phosphatases alter the phosphorylation state of synapsins, including MAP kinase^50–52^, CdK5^53–55^, CaMKII^51,52,56,57^ and PP2A/PP1^51,56^ (**Figure S5**). Out of all these molecules, CaMKII is expressed at very low levels in INs ^58,59^, and although MAP kinase can be activated by Ca^2+^, its increased activity would not explain the reduced *RRP* in 2.5 mM Ca^2+^. CdK5 and PP2A/PP1 are not modulated by Ca^2+^. By contrast, the protein phosphatase PP2B (i.e., calcineurin) is a ubiquitous Ser/Thr phosphatase activated by elevated Ca^2+^ levels and subsequent activation of calmodulin ^60^. Calcineurin is known to dephosphorylate synapsins and antagonize the effect of MAP kinase and CdK5 on these molecules ^51,61^. Therefore, if the reduced *RRP* in [𝐶𝑎]^2+^=2.5 mM is due to synapsins dephosphorylation by calcineurin, we would expect cyclosporin A (1 µM), an inhibitor of calcineurin, to increase the *RRP* and rescue its modulation by DHK in 2.5 mM Ca^2+^. To test this hypothesis, we repeated our single and train stimulation in slices incubated with cyclosporin A for 30-60 min (**Figure 4O-U**). Consistent with our hypothesis, DHK reduced the IPSC amplitude (***p=3.7e-4; **Figure 4O-Q**) and the *RRP* in [𝐶𝑎^2+^]_𝑜_=2.5 mM in the presence of cyclosporin A (*p=0.011; **Figure 4R-U**). The findings that *(i)* DHK reduced the IPSC amplitude and *RRP* in slices maintained in cyclosporin A and [𝐶𝑎^2+^]_𝑜_=2.5 mM (**Figure 4o-u**) and that *(ii)* the effect of DHK was no longer detectable in slices maintained in cyclosporin A and MSOP in [𝐶𝑎^2+^]_𝑜_=2.5 mM (**Figure 4V-BB**), indicate that the inability to detect an mGluRIII- dependent reduction of inhibition onto CA1-PCs in [𝐶𝑎^2+^]_𝑜_=2.5 mM is likely due to an altered balance in the phosphorylation state of synapsins mediated by calcineurin.

### mGluRIII receptors reduce proximal and distal inhibition onto CA1-PCs

The EM data in **Figure 1** show that mGluRIII is expressed at proximal and distal I-synapses onto CA1-PCs, which are mostly formed by PV- and SST-INs, respectively. To selectively activate I-afferents from PV- or SST-INs and determine whether these largely non-overlapping inputs are equally susceptible to glutamate uptake blockade, we performed stereotaxic injections of a flexed viral construct encoding ChR2 in *Pvalb^Cre/+^* and *Sst^Cre/+^* mice (**Figure 5A**). We then recorded optically evoked IPSCs (oIPSCs) in CA1-PCs. DHK reduced the amplitude of oIPSCs from PV-INs to 74±8% (n=10; *p=0.011; **Figure 5B-E**), and that of oIPSCs from SST-INs to 74±4% (n=10; ***p=7.9e-5; **Figure 5F-I**). Given the different spatial distribution of PV- and SST-inputs onto the dendritic tree of CA1-PCs (**Figure 5J**), inhibition from these two different classes of INs is prone to different levels of electrotonic attenuation, as the synaptic currents propagate towards the soma (**Figure S7**). Accordingly, cable theory suggests that with somatic voltage-clamp recordings, fast currents arising at a distance from the soma are attenuated more than currents with slower kinetics arising in more proximal compartments ^62^. This would suggest that glutamate spillover could induce a larger reduction of distal compared to proximal inhibition. We tested this hypothesis using a compartmental model that reproduced not only the realistic morphology of CA1-PCs but also the voltage escape associated with somatic voltage-clamp recordings^63,64^ (**Figure S7A-D**). We constrained the synaptic weight of PV- inputs in the model (*w_PV_=*485 pS) to ensure that it would generate somatic mIPSCs with similar amplitude and kinetics compared to those recorded experimentally (**Figure 3A-D**; **Figure S7E-F**). The synaptic weight of SST- inputs was set to be the same (*w_PV_*=*w_SST_*=485 pS; **Figure S7E-F**). The release probability of PV-INs was set to *Pr_PV_*=0.10 in [𝐶𝑎^2+^]_𝑜_=1.2 mM, a value we derived using a power-law relationship from a starting value of *Pr_PV_*=0.8 measured in [𝐶𝑎^2+^]_𝑜_=2.0 mM ^19,65,66^ (**Figure S7G**). The model showed that evoking GABA release from 4 out of 40 PV inputs generated somatic oIPSCs with the same amplitude and kinetics of the PV-oIPSCs recorded experimentally (**Figure 5K**, *top*). Based on 3D EM reconstructions, the number of SST-inputs onto CA1-PCs is 8.3 times larger than that of PV-INs (i.e., 40·8.3=332) ^67^. Assuming that all these terminals are activated synchronously by the full-field light stimulus in our optogenetics experiments and given that oIPSCs similar to those detected experimentally could be generated by evoking release from 18 SST-inputs in the model (**Figure 5K**, *bottom*), we estimated the release probability of SST-INs to be *Pr_SST_*=18/332=0.05 (**Figure S7G**). Based on the data shown in **Figure 3**, promoting glutamate spillover reduces the size of the *RRP*, which is composed of *N* release sites. Our simulations showed that a 25-28% reduction in the amplitude of somatically-recorded PV- or SST-oIPSCs (**Figure 5E, I**) could be obtained by reducing the total number of PV release sites from 40 to 30, and that of SST-INs from 332 to 260. This is analogous to reducing the number of active release sites of PV-INs from 4 to 3 and of SST-INs from 18 to 13, representing a 25% and 28% reduction, respectively. Therefore, experimental conditions that promote glutamate spillover and mGluRIII activation induce minimal – if any – change in the relative strength of proximal versus distal inhibition onto CA1-PCs (**Figure 5L**). The relevance of this effect in the context of intact networks and encoding of spatial information in CA1-PCs is analyzed below.

**Figure 5.**
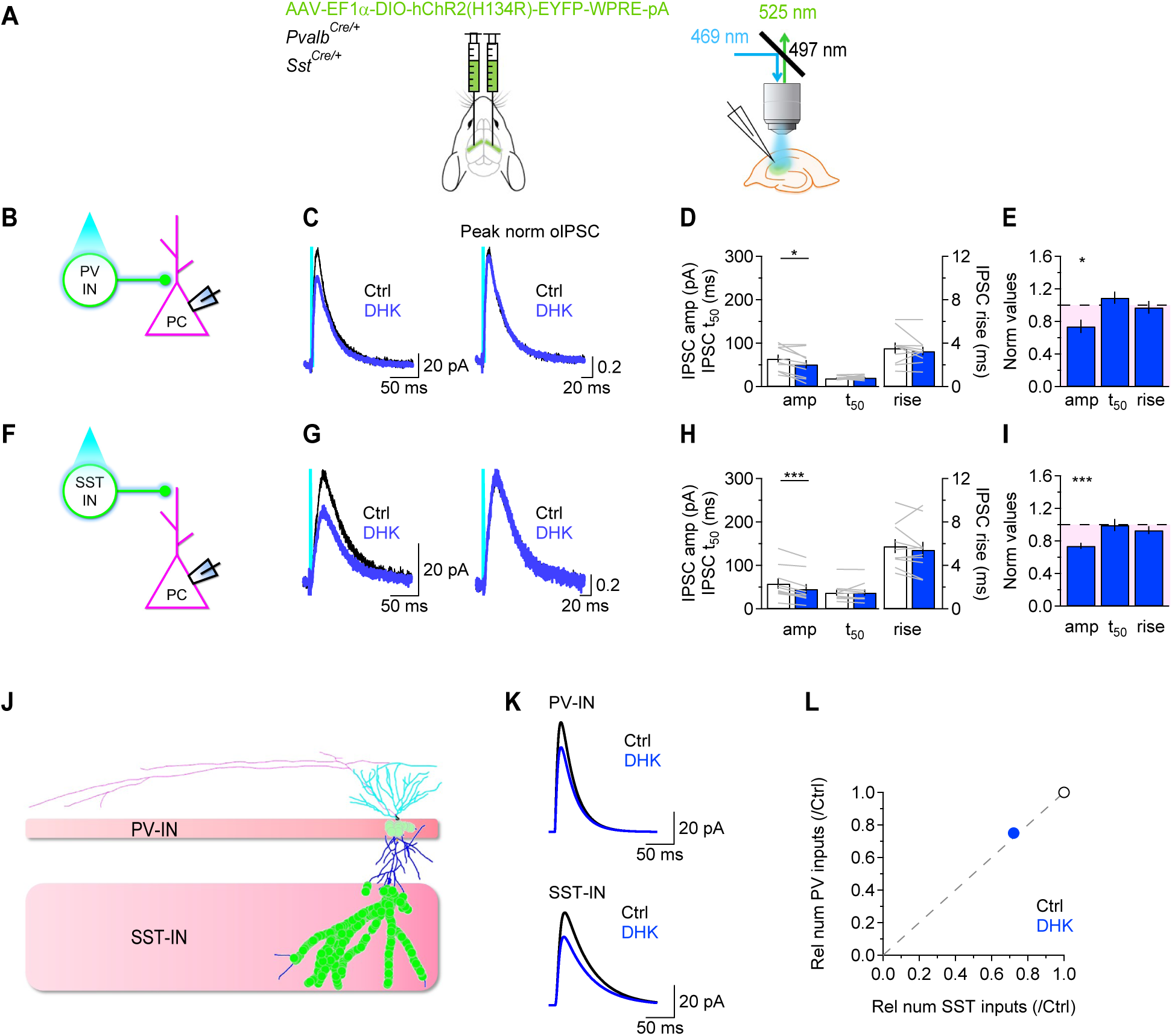
Proximal and distal inhibition onto CA1-PCs are modulated by mGluRIII. **(A)** Schematic representation of the stereotaxic injections of viral constructs encoding ChR2 in hippocampal area CA3 of *Pvalb^Cre/+^* and *Sst^Cre/+^* mice, and of the experiment settings. **(B)** Scheme describing optogenetic activation of PV-INs. **(C)** oIPSCs evoked by optical stimulation of PV-INs before (*black*) and after DHK (100 µM; *blue*). The traces on the right are normalized by the iIPSC peak. **(D)** Summary of the effect of DHK on the oIPSC amplitude and kinetics, in *Pvalb^Cre/+^* mice. **(E)** Summary of the effect of DHK on the oIPSC amplitude and kinetics, normalized by their values in control conditions. **(F-I)** As in B-E, for oIPSCs evoked by optical stimulation of SST-INs. **(J**) Morphology of a CA1- PCs receiving PV-inputs onto the soma and the proximal portion of the apical dendrite (0-50 µm; *light green dots*), or SST-inputs onto the distal portion of the apical dendrite (>200 µm; *green dots*). **(K)** Example of oIPSCs generated with a compartmental model of a CA1-PCs in response to activation of PV- (*top*) or SST-inputs (*bottom*). The model reproduces the effect of DHK on the amplitude and kinetics of these events detected experimentally (C, G). **(L)** A 25-28% reduction in the amplitude of the oIPSC recorded at the soma can be obtained by reducing the number of active PV-inputs from 4 to 3, and that of SST-inputs from 11 to 8. The scatter plot shows these results as relative effects in control conditions (*black*) and in DHK (*blue*).

### mGluRIII receptors increase the accuracy of space representation by CA1-PCs

The *in vitro* experiments described above indicate that mGluRIII modulates synaptic transmission in area CA1, limiting excitation and inhibition onto CA1-INs and reducing inhibition, but not excitation, onto CA1-PCs under physiological [𝐶𝑎^2+^]_𝑜_conditions. Since CA1-PCs are crucial for encoding spatial representations, and the influence of mGluRIII on these representations has not been previously explored, we conducted a series of *in vivo* experiments to investigate these phenomena. In these experiments, we performed simultaneous whole-cell patch-clamp recordings from CA1-PCs using a biocytin-containing internal solution and extracellular local field potential (LFP) recordings in area CA1 (**Figure 6A**, *top*). The glass electrode used to record the local field potential also contained either the mGluRIII antagonist MSOP (100 mM) or a vehicle solution. *Post hoc* analysis of the solution’s spatial spread (∼500 µm) confirmed the efficacy of the local MSOP or vehicle delivery (**Figure 6A**, *bottom*). In awake and anesthetized mice, we found that mGluRIII activation occurred even without GLT-1 blockade, likely due to the higher levels of ongoing synaptic activity *in vivo* (**Figure 6, S8**). In anesthetized mice, the mGluRIII antagonist MSOP caused a constant depolarization of the membrane potential (V_m_) (n=10; **Figure S8A-D**) and an increase in the AP firing frequency of CA1-PCs (**Figure S8E-F**). These effects were not artifacts of the pressure application method used to deliver MSOP, as the control experiments with vehicle solution showed no changes in V_m_ or firing frequency (n=7; **Figure S8G-L**). Thus, mGluRIII activation in anesthetized mice reduces the firing output of CA1- PCs.

**Figure 6.**
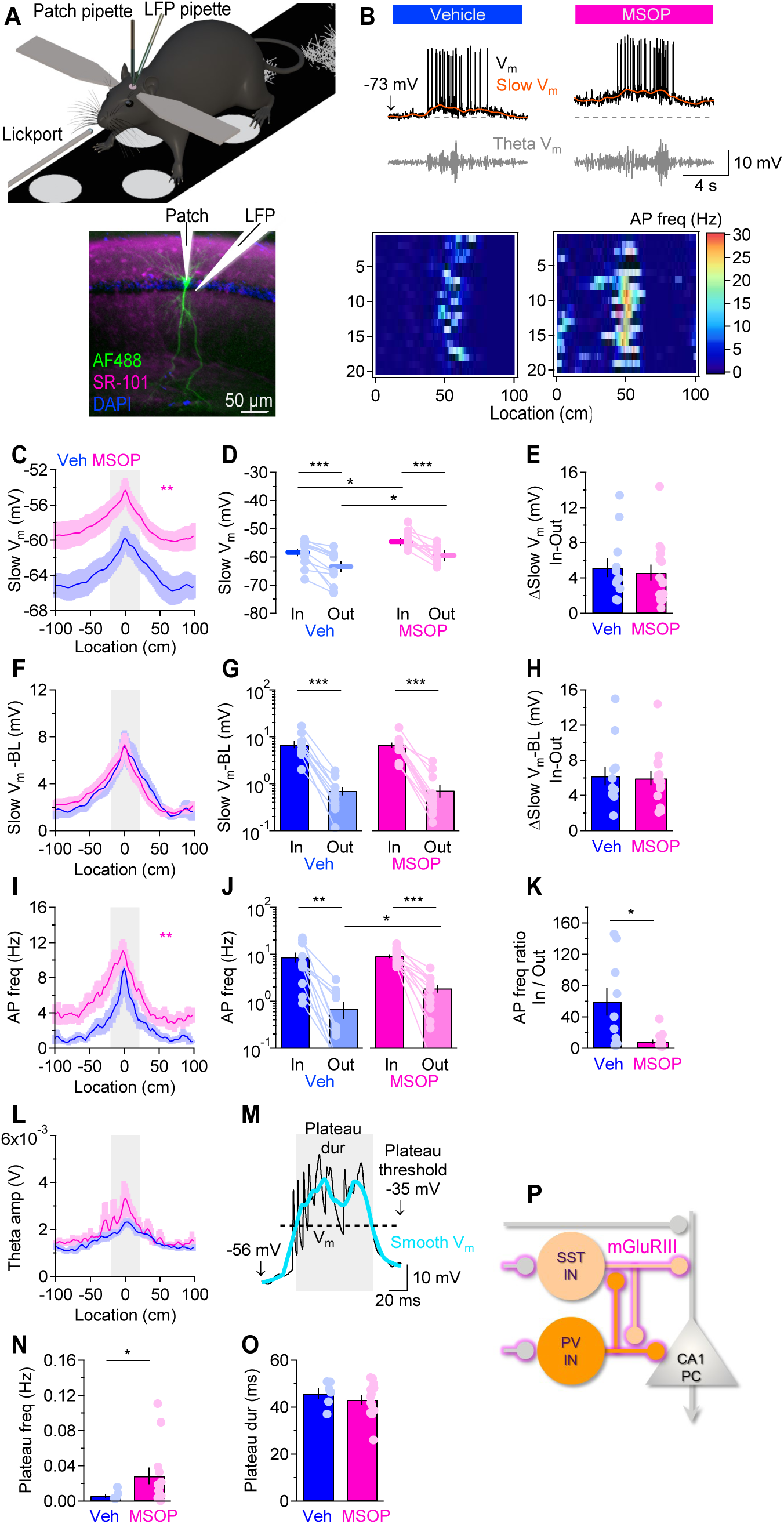
Spillover-mediated mGluRIII activation reduces place field formation and strength. **(A)** *Top*, Schematic of the linear treadmill and recording apparatus for the *in vivo* electrophysiology experiments. *Bottom*, Confocal image of biocytin-filled CA1-PCs (*green*), SR-101 labeling to depict the area affected by MSOP/vehicle (*magenta*), and DAPI (*blue*) in a section of the mouse hippocampus. **(B)** Example of raw V_m_ (*black*), low-pass filtered V_m_ (slow V_m_; *orange*), and theta-filtered intracellular V_m_ (*gray*) for CA1-PC place cells recorded in vehicle and MSOP. The heatmaps represent the spatial firing rate of these cells shown for 20 consecutive laps. The dashed lines marks -73 mV. **(C)** Slow *V*_m_ ramp of CA1 place cells. The gray shaded area represents the location of the in-field region. **(D-E)** Summary of the in-field, out-of-field, and the difference in the in-field vs. out-of-field absolute slow V_m_ values. MSOP depolarizes the in-field and out-of-field slow V_m_ by ∼5mV. **(F)** Baseline (BL) V_m_-subtracted slow V_m_, showing no change in the absolute amplitude of the slow V_m_ ramp depolarization between vehicle and MSOP. **(G-H)** Summary of the baseline subtracted in-field, out-of-field, and the difference in the in-field vs. out-of-field baseline-subtracted slow V_m_ values. MSOP does not change the amplitude of the baseline-subtracted slow V_m_. **(I)** Spatial dependence of AP frequency. **(J-K)** Summary of the in-field, out-of-field, and in-field vs. out-of-field AP frequency ratio. MSOP decreases the in-field to out-of-field AP frequency ratio. **(L)** Spatial dependence of the amplitude of the intracellular theta activity. **(M)** Example of raw V_m_ (*black*) and smoothed, subthreshold V_m_ (*cyan*) during a spontaneous plateau. The gray shaded area represents the duration of the plateau, calculated from the threshold value of -35 mV. **(N-O)** Summary graphs of the plateau frequency and duration, respectively. MSOP increased the plateau frequency without altering the plateau duration. **(P)** Diagram of hippocampal connectivity and synaptic location of mGluRIII-dependent modulation.

To examine how mGluRIII shapes the activity of CA1 place cells in awake-behaving mice, we recorded next from CA1 place cells in mice trained to run on a linear treadmill for a sucrose reward (n=12 in vehicle; n=15 in MSOP; **Figure 6A**). The recordings included both pre-existing place cells (n=4 in vehicle; n=9 in MSOP) and silent cells, which were experimentally induced to form place fields using the behavioral time scale plasticity (BTSP) protocol (n=8 in vehicle; n=6 in MSOP)^68^. The intracellular V_m_ dynamics were decomposed into three components: suprathreshold AP firing, a slow subthreshold V_m_ component (<2 Hz), and an intracellular theta oscillation component (5-10 Hz; **Figure 6B**). Consistent with prior work ^69^, vehicle-treated control CA1 place cells showed a ramp-like depolarization in the slow V_m_, a place-specific increase in AP firing rate, and increased theta oscillation amplitude (**Figure 6B**). Compared to the control cell population, CA1 place cells in the MSOP-treated group exhibited a ∼5 mV position-independent depolarizing shift in the slow V_m_ (**Figure 6C-H**), leading to a significantly increased firing rate out-of-field (**Figure 6I-J**). The ratio of in-field to out-of-field firing decreased (**Figure 6K**), suggesting that mGluRIII activation contributes to maintaining spatial selectivity. The intracellular theta oscillation amplitude remained unchanged (**Figure 6L**). To further explore these effects, we analyzed complex spike bursts, or “plateaus”, which are mediated by dendritic plateau potentials, large voltage events initiated in the distal dendrites of CA1-PC ^68,70,71^ (**Figure 6M-O**). MSOP application increased plateau frequency but did not affect plateau duration. If mGluRIII activation *in vivo* is driven by glutamate spillover, we would anticipate that the effects of MSOP would closely mirror those observed when extracellular diffusion is reduced using dextran ^72,73^. Consistent with this hypothesis, dextran produced similar effects on V_m_ dynamics and spatial selectivity as MSOP (n=13 in dextran; **Figure S9**). Together, these results suggest that spillover-mediated mGluRIII activation is critical for encoding spatial information in CA1 place cells. The effects we observed are consistent with mGluRIII activation causing a net increase in synaptic inhibition of CA1 place cells *in vivo*.

## Discussion

The main finding in this work is that, under experimental conditions that closely match the physiological Ca^2+^ concentration in the cerebrospinal fluid ^18^ and promote mGluRIII activation, there is no target cell-specific control of inhibition onto CA1-PCs. The same experimental conditions, however, lead to reduced excitation onto CA1-INs but not PCs. These results differ, in part, from those collected in slice physiology experiments performed using a higher extracellular [𝐶𝑎^2+^]_𝑜_, which showed that mGluRIII reduces excitation and inhibition onto CA1-INs, not CA1-PCs ^3,16,22,74^. We identify PKA-dependent phosphorylation of synapsins as a key mechanism that limits the inhibition of CA1-PCs by reducing the size of the RRP. A key question is how increasing [𝐶𝑎^2+^]_𝑜_ prevents the mGluRIII- dependent modulation of synaptic inhibition of CA1-PCs. Our experimental data show that varying [𝐶𝑎^2+^]_𝑜_ alters the activation of the Ca^2+^-dependent phosphatase calcineurin, which can dephosphorylate synapsins. We speculate that, at high [𝐶𝑎^2+^]_𝑜_, synapsins may be in a net dephosphorylated state due to the presence of high affinity, Ca^2+^- dependent phosphatases in the bouton ^75^. Therefore, we propose that varying [𝐶𝑎]^2+^ can modify the mGluRIII- dependent control of inhibition onto CA1-PCs by altering the phosphorylation state of proteins like synapsins, which are involved in regulating the *RRP*.

We do not know why the mGluRIII-dependent modulation of inhibition onto CA1-PCs is sensitive to [𝐶𝑎^2+^]_𝑜_ but excitation is not. We acknowledge that synapsins have unique functions at excitatory versus inhibitory synapses. At excitatory synapses, synapsins maintain the reserve pool of glutamatergic vesicles, whereas at inhibitory synapses they regulate the size of the RRP ^76^. However, there is currently no indication that the biophysical properties of these proteins differ across pre-synaptic terminals onto different cell types, making this an unlikely scenario. Our ultrastructural analysis shows that one mGluRIII type – mGluR7a – is more abundant at I- compared to E-terminals, but we do not know whether target cell-specific differences in the localization of mGluRIII relative to the RRP or to other proteins implicated with the complex regulatory pathway that controls the RRP size exist. These ultrastructural features, as well as lower levels of cross-talk between E compared to E-to-I synapses, and different calcium buffering capacities of E/I synapses could all contribute to the higher [𝐶𝑎^2+^]_𝑜_-sensitivity of inhibition compared to excitation onto CA1-PCs ^77^. There is, in fact, anatomical data showing that, in the hippocampus, the distance between adjacent glutamatergic synapses is twice as large as that between glutamatergic and GABAergic synapses (∼767 nm versus ∼69 nm, respectively) ^78^. Target cell-specific differences in the ability of activity-dependent retrograde signals to control GABA release could make the scenario even more complex ^79,80^.

The finding that the effects of mGluRIII activation are mediated by PKA inhibition and dephosphorylation of its vesicle-associated target proteins, synapsins, is reminiscent of the signaling pathways that regulate the occupancy of the *RRP* at E-synapses in the cerebellum, suggesting that this regulatory pathway may be widespread across E/I synapses across the brain ^37^. Our data show that an mGluRIII-dependent control of inhibition onto principal cells occurs in SS1. An essential finding of this work, however, is that it identifies mGluRIII as a family of receptors capable of inhibiting the PKA-synapsins signaling pathway, leading to a reduction in *RRP*. PKA may not be the only protein implicated with regulating the phosphorylation state of synapsins, as there are multiple protein kinases and phosphatases known to control the phosphorylation state of these molecules ^50–57^. However, the fact that blocking PKA activity prevents the reduction of inhibition onto CA1-PCs in the presence of DHK suggests PKA is a key player in this regulatory mechanism.

The inability to detect an mGluRIII-dependent modulation of GABA release onto CA1-PCs in the absence of glutamate transporter blockers *in vitro* but not *in vivo* is consistent with different levels of ongoing synaptic activity in living organisms and with the fact that many glutamatergic afferents to the hippocampus are resected during slice preparation. This finding also implies that mGluRIII may control the excitability of CA1-PCs in an activity-dependent manner, reducing it at increasing activity levels of excitatory networks. In this case, one could ask whether there is any potential therapeutic benefit of using mGluRIII antagonists to dampen epileptic seizures or cognitive dysfunction in diseases associated with increased extracellular glutamate levels, like Alzheimer’s disease. A reduced expression of mGluRII/III has been detected in human Ammon’s horn sclerosis ^81^ and the pilocarpine-induced *status epilepticus* model of chronic epilepsy in rodents ^82–85^. However, the use of mGluRIII agonists has shown mixed responses in animal models of epilepsy. Some groups reported that the mGluRIII agonists, L-AP4 and L-SOP, are proconvulsive and mGluRIII antagonists like MCPA are anticonvulsive ^86^. Others showed that mGluRIII agonists are anticonvulsants in generalized motor and focal seizures ^87–96^ and that other antagonist, such as MAP4 and MPPG, are proconvulsant ^86,89^. A probable reason for the discrepancy of these results may lie in the limited specificity of the pharmacological tools used to modulate mGluRIII receptors and in the heterogeneous mGluRIII expression and recruitment at different synapses throughout the hippocampal circuitry and in different epilepsy models, making the results particularly challenging to interpret.

The *in vivo* experiments underscore a pivotal role for mGluRIII-mediated synaptic modulation in shaping spatial representations in hippocampal area CA1 and significantly extend our current understanding of the role of mGluRIII for behaviorally relevant neural circuit dynamics. The impact of mGluRIII modulation was evident even without pharmacological manipulations, such as glutamate uptake blockade, indicating that the *in vivo* environment provides sufficient glutamate spillover to activate these receptors. This highlights the complementary role of studying synaptic transmission in slices and within intact networks, where ongoing neural activity reveals physiological mechanisms that may be obscured in reduced preparations.

We observed that mGluRIII activation has effects consistent with an overall increased level of synaptic inhibition, thus altering the spatially tuned firing patterns of individual CA1-PCs and enhancing the CA1 network’s ability to sparsely encode spatial information. Our findings align with prior *in vivo* work highlighting the critical role of inhibition in maintaining flexible and selective coding of CA1 place cells while adding to these results by demonstrating that mGluRIII activation plays a role in modulating inhibition ^97–99^. Specifically, mGluRIII activation appears to also affect distal GABAergic inhibition onto CA1-PCs, as evidenced by the increased frequency of dendritically driven plateaus following the mGluRIII blockade.

Based on our *in vitro* recordings, we hypothesize that the observed mGluRIII-mediated increase in net synaptic inhibition of CA1-PCs is the result of a combination of reduced excitation of disinhibitory CA1-INs and reduced activation of IN-to-IN synapses. Both effects would enhance PV- and SST-IN activity, thereby exerting a net suppressive effect on CA1-PC activity *in vivo*. The potential behavioral implications of these findings are profound. By modulating the E/I ratio in the CA1 circuit, mGluRIII would be well-positioned to influence the accuracy and stability of spatial representations in CA1. This modulation may play a critical role in adaptive navigation and spatial memory formation, potentially enabling dynamic adjustments to the E/I balance necessary to encode new environments while preserving established spatial maps. Such flexibility is thought to be essential for effective learning and memory processes ^100^, underscoring the functional relevance of mGluRIII receptors in hippocampal circuit dynamics.

## Resource availability

All primary data used in this work and complete statistical analyses for each figure have been deposited in an Open Science Framework repository (https://osf.io/ucj2d/). All files pertaining to the NEURON simulations were uploaded to the ModelDB database (https://senselab.med.yale.edu/modeldb/, ModelDB acc.n. TBD).

## Acknowledgements

We would like to thank G.C. Todd and P.A. Albrecht for mouse colony management and genotyping. Special thanks to B. L. Sabatini for valuable discussions.

## Authors contributions

Conceptualization: CG, FC, AS Data curation: MAP, MM, ND, CG, AS Formal analysis: MAP, MM, ND, AS Funding acquisition: CG, FC, AS Investigation: MAP, MM, TJR, AS Methodology: ND, CG, AS Project administration: AS Resources: CG, FC, AS Software: MAP, ND, CG, AS Supervision: CG, FC, AS Validation: MM, AS Visualization: MAP, MM, AS Writing – original draft: MM, CG, FC, AS Writing – review & editing: CG, FC, AS

## STAR ★ Methods

### Experimental model and study participant details

#### Mice

All experimental procedures were performed in accordance with the guidelines of the National Institutes of Health and were approved by the Institutional Animal Care and Use Committee at the State University of New York (SUNY) Albany, Brandeis University and guidelines described in the National Institutes of Health’s Guide for the Care and Use of Laboratory Animals. Unless otherwise stated, all mice were group housed and kept under 12 h:12 h light:dark conditions. Food and water were available *ad libitum* to all mice throughout the 24-hour period. For our experiments, we used mice of either sex from the mouse lines reported below. *In vitro* electrophysiology experiments were performed in 4-8 week old mice. *In vivo* electrophysiology experiments were performed in 8-14 week old mice.

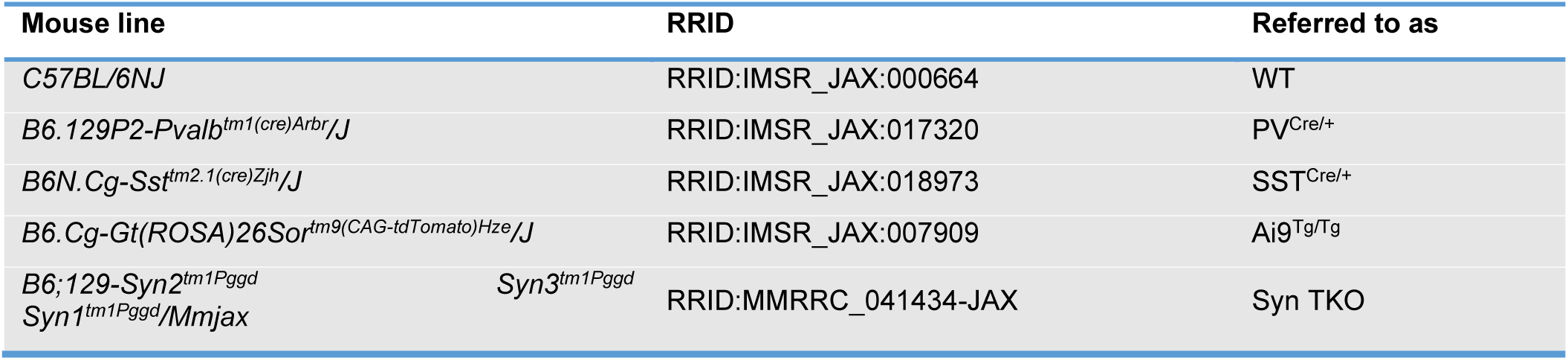

Genotyping was performed on toe tissue samples of P7–P10 mice. Briefly, tissue samples were digested at 55°C overnight with shaking at 330 rpm in a lysis buffer containing the following (in mM): 100 Tris-HCl base, pH 8, 5 EDTA, and 200 NaCl, along with 0.2% SDS and 100 mg/ml proteinase K. Following heat inactivation of Proteinase K at 97°C for 10 min, DNA samples were diluted 1:1 with nuclease-free water. The PCR primers were purchased from Eurofins Genomics and their nucleotide sequences are listed below.

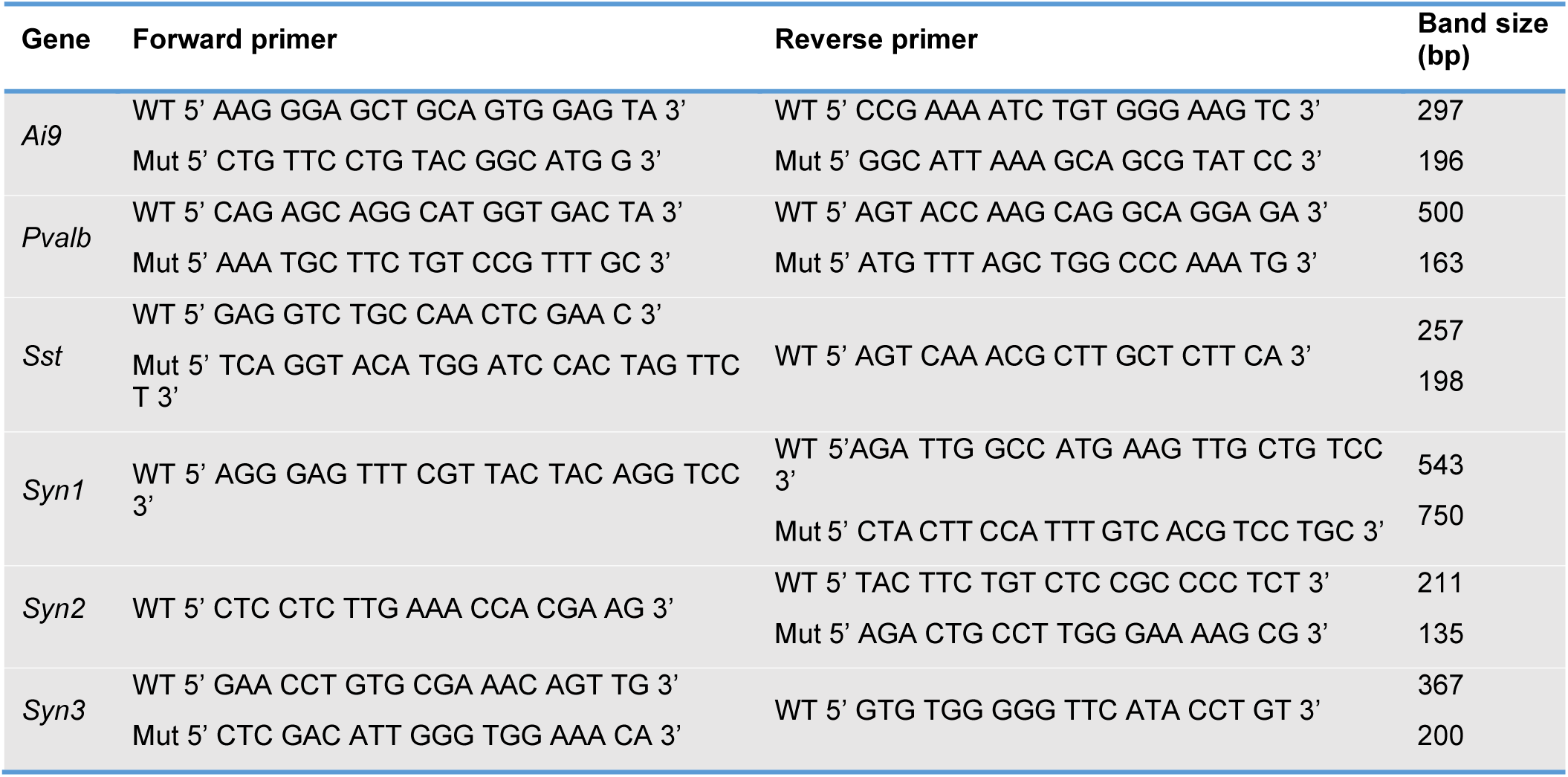

PCR was carried out using the KAPA HiFi Hot Start Ready Mix PCR Kit (KAPA Biosystems, KK2602). Briefly, 12.5 µl of 2× KAPA HiFi Hot Start Ready Mix was added to 11.5 µl of a diluted primer mix (0.5-0.75 mM final concentration for each primer) and 1 µl of diluted DNA. The PCR cycling protocols are described in the table below.

| ***Ai9, Pvalb,***<br>***Sst, Syn2*** | **Initiation/melting** | **Denaturation** | **Annealing** | **Elongation** | **Amplification** | **Hold** |
| --- | --- | --- | --- | --- | --- | --- |
| *Temperature (°C)* | 95 | 95 | 60 | 72 | 72 | 4 |
| *Duration (min)* | 3 | 0.25 | 0.25 | 0.25 | 1 | ∞ |
| *Cycles* | 1 | 35 |  |  | 1 |  |

| ***Syn1*** | **Initiation/melting** | **Denaturation** | **Annealing** | **Elongation** | **Amplification** | **Hold** |
| --- | --- | --- | --- | --- | --- | --- |
| *Temperature (°C)* | 95 | 95 | 56 | 72 | 72 | 4 |
| *Duration (min)* | 3 | 0.25 | 0.25 | 0.5 | 1 | ∞ |
| *Cycles* | 1 | 35 |  |  | 1 |  |

| ***Syn3*** | **Initiation/melting** | **Denaturation** | **Annealing** | **Elongation** | **Amplification** | **Hold** |
| --- | --- | --- | --- | --- | --- | --- |
| *Temperature (°C)* | 95 | 95 | 64 | 72 | 72 | 4 |
| *Duration (min)* | 3 | 0.25 | 0.25 | 0.25 | 1 | ∞ |
| *Cycles* | 1 | 35 |  |  | 1 |  |

### Method details

#### Stereotaxic intracranial AAV injections

AAV5-EF1a-DIO-hChR2(H134R)-EYFP-WPRE-pA (University of North Carolina Vector Core, Chapel Hill, NC) was injected into PV^Cre/+^ and SST^Cre/+^ mice. Male and female mice (P14-16) were anesthetized with isoflurane (induction: 5% in 100% O_2_ at 1-2 l/min; maintenance: 3% in 100% O_2_ at 1-2 l/min) and placed in the stereotaxic frame of a motorized drill and injection robot (Neurostar GmbH, Tübingen, Germany). After making a skin incision and thinning the skull under aseptic conditions, we injected 200 nl of the viral constructs bilaterally in *stratum radiatum* using a Hamilton syringe at a rate of 50 nl/min. The injection coordinates from bregma were AP: -1.9 mm, ML: ±1.6 mm, DV: -1.4 mm. After the stereotaxic injections, the mice were returned to their home cage and used for slice physiology experiments 2-6 weeks after surgery.

#### Acute slice preparation

Acute coronal slices of the mouse hippocampus were obtained from C57BL/6J mice of either sex (4-8 week old), deeply anesthetized with isoflurane and decapitated in accordance with SUNY Albany Animal Care and Use Committee guidelines. The brain was rapidly removed and placed in ice-cold slicing solution bubbled with 95%O2/5%CO2 containing the following (in mM): 119 NaCl, 2.5 KCl, 0.5 CaCl_2_, 1.3 MgSO_4_·H_2_O, 4 MgCl_2_, 26.2 NaHCO_3_, 1 NaH_2_PO_4_, and 22 glucose, 320 mOsm, pH 7.4. The slices (250 µm thick) were prepared using a vibrating blade microtome (VT1200S; Leica Microsystems, Buffalo Grove, IL). Once prepared, the slices were stored in slicing solution in a submersion chamber at 36°C for 30 min and at RT for at least 30 min and up to 5 hours.

#### In vitro electrophysiology recordings and optogenetics

Unless otherwise stated, the recording solution contained the following (in mM): 119 NaCl, 2.5 KCl, 1.2 CaCl_2_, 1 MgCl_2_, 26.2 NaHCO_3_, and 1 NaH_2_PO_4_, 22 glucose, 300 mOsm, pH 7.4. We identified the hippocampus under bright field illumination using an upright fixed-stage microscope (BX51 WI; Olympus Corporation, Center Valley, PA). To record electrically evoked currents, we delivered constant voltage stimuli (50-100 ms) to a bipolar stainless-steel electrode (Cat# MX21AES(JD3); Frederick Haer Corporation, Bowdoin, ME) positioned in *stratum radiatum*, ∼100 µm away from the recorded cell. To activate ChR2-expressing fibers from PV- and SST-INs, we used 5 ms-long light pulses generated by a SOLA-SE light engine (Lumencor, Beaverton, OR) and filtered using a green FITC filter set (469/497/525 nm). The light power at the sample plane was ∼250 µW and the light pulses were delivered at intervals of 30 s using whole-field illumination through a 40× water immersion objective (LUMPLFLN40XW; Olympus, Center Valley, PA). When testing the effect of PKA inhibition, we incubated the hippocampal slices with KT5720 (1 µM) for 30-60 min prior to recording. Whole-cell, voltage-clamp patch-clamp recordings were made using patch pipettes containing (in mM): 120 CsCH_3_SO_3_, 10 EGTA, 20 HEPES, 2 MgATP, 0.2 NaGTP, 5 QX-314Br, 290 mOsm, pH 7.2. All recordings were obtained using a Multiclamp 700B amplifier (Molecular Devices, San Jose, CA) and filtered at 10 KHz, converted with an 18-bit 200 kHz A/D board (HEKA Instrument, Holliston, MA), digitized at 10 KHz, and analyzed offline with custom-made software (A. S.) written in IgorPro 6.37 (Wavemetrics, Lake Oswego, OR). Patch electrodes (#0010 glass; Harvard Apparatus, Holliston, MA) had tip resistances of ∼5 MΩ. Series resistance (∼20 MΩ) was not compensated but was continuously monitored and experiments were discarded if this changed by >20%. The series resistance was subtracted from the total resistance to estimate the cell membrane resistance of neurons and astrocytes. Extracellular recordings were performed using electrodes with a tip resistance of 2-5MΩ, filled with the recording solution described above. The electrode resistance was continuously monitored, and experiments were discarded if this changed by >20%. Tetrodotoxin (TTX), 2,3-Dioxo- 6-nitro-1,2,3,4-tetrahydrobenzo[f]quinoxaline-7-sulfonamide disodium salt (NBQX) and D,L-2-Amino-5- phosphonopentanoic acid (APV) were purchased from Hello Bio (Princeton, NJ; Cat# HB1035, HB0443, HB0251, respectively). (3S)-3-[[3-[[4-(Trifluoromethyl)benzoyl]amino]phenyl]methoxy]-L-aspartic acid (T-TBOA) was purchased from Tocris (Minneapolis, MN; Cat# 2532). All other chemicals were purchased from Millipore Sigma (Burlington, MA). All recordings were performed at RT.

#### Surgeries for in vivo electrophysiology experiments and training

All mice were housed under a reversed 12-h light/12-h dark cycle. All surgical procedures were performed under deep isoflurane anesthesia (4% induction, 2-3% maintenance). After locally applying topical anesthetics, the scalp was removed and cleaned. Then the skull was leveled, and locations for the craniotomies for the whole-cell patch clamp and LFP recordings were marked using stereotactic coordinates (whole-cell recordings: AP -2.0 mm, ML ±1.7 mm; LFP recordings: AP -3.55 mm, ML ±1.7 mm). Then, the skull was covered with cyanoacrylate glue (Loctite). As soon as the glue had dried, custom-made titanium head bars with an opening over the hippocampus were attached to the skull using dental acrylic (Ortho-Jet, Lang Dental). Following at least a week of recovery, mice were placed on water restriction (1.5 ml/day). After at least 3 days of water restriction mice were handled by the experimenter for at least five days (15 min/day). Next, mice were trained to run on the treadmill for a 10% sucrose (in water) reward delivered through a licking port once for every complete rotation of the belt, i.e., a lap of 200 cm. Mice were supplemented with additional water after training sessions to guarantee a daily water intake of 1.5 ml/day. Mice were trained until the number of laps completed per day was >100 laps/hr and was comparable for 2-3 days. The typical treadmill training duration was 4-7 days. The day before the first recording session, two small craniotomies (∅ = 0.5 mm), one for the whole-cell patch and one for the LFP electrode, were drilled at the previously marked positions under isoflurane anesthesia. The dura was, if possible, left intact, and the craniotomies were covered with a biocompatible silicone elastomer (KWICK-CAST, World Precision Instrument, Sarasota, FL).

#### Behavioral setup

The linear track treadmill consisted of a belt made from velvet fabric (McMaster Carr) enriched with visual and tactile cues ^68,97,101^. The belt was self-propelled and divided up into three sectors, each with a different type of local cue. The sucrose/water reward (5-7 µl) was delivered through a custom-made lick port controlled by a solenoid valve (Lee Company, Westbrook, CT), at a point located 130 cm from the beginning of the belt. A single photoelectric sensor (Sparkfun, Niwot, CO) was used to reset the location measurement in each lap (=lap start). The mouse running speed was measured using an encoder attached to one of the wheel axles (Cat# HEDS– 5540#I06, Avago, San Jose, CA). The valve, sensor, and encoder were controlled with microprocessor-controlled behavioral control system (Bpod, Sanworks, Rochester, NY). A separate Arduino Teensy 3.5 microprocessor interfaced with a custom-made MATLAB GUI was used to control the current injections based on the position of the mouse on the belt.

#### In vivo electrophysiology

All whole-cell recordings were made in current clamp mode using a model 2400 patch-clamp amplifier (A-M Systems, Sequim, WA). The LFP recordings were conducted using either the model 2400 patch-clamp amplifier or a model 1700 differential AC amplifier (A-M Systems, Sequim, WA). All signals were digitized at 20 kHz using the National Instruments PCIs-6341 card and the open-access MATLAB-based software (WaveSurfer, Janelia Research Campus). Liquid junction potentials were not corrected. CA1-PC recordings were either aborted or excluded from further analysis if access resistance exceeded a predefined threshold of 50 MW.

For each mouse, the first step involved determining the depth of the CA1 pyramidal cell (CA1-PC) layer using the LFP signals. Glass electrodes (1.5-3.5 MΩ), filled with 0.9% NaCl, were mounted vertically on the micromanipulator to be used later for the whole-cell recordings (Luigs and Neumann, Ratingen, Germany) and slowly advanced toward the hippocampus. The LFP signal was monitored via an audio amplifier (Grass Technologies), and the CA1- PC layer, typically located 1.1–1.3 mm below the brain surface, was identified by the presence of theta-modulated spikes and increased ripple amplitude. The extracellular electrode to be used for the simultaneous LFP recordings was then mounted on a second micromanipulator (Narishige, Amityville, NY) with a 40-45° angle relative to bregma. This electrode was advanced through the second, more posterior craniotomy until the CA1-PC layer was identified using the same criteria. Two LFP signals were recorded simultaneously to measure the phase shift between the LFP electrode position and the site of whole-cell patch recording. Subsequently, the vertically oriented LFP electrode was replaced with whole-cell patch electrodes (8-12 MΩ) filled with intracellular solution containing (in mM): 134 K-gluconate, 6 KCl, 10 HEPES, 4 NaCl, 0.3 MgGTP, 4 MgATP, and 14 Tris-phosphocreatine. Biocytin (0.2%) was added to the intracellular solution. Whole-cell electrodes were vertically advanced through the cortex with 8-9 psi of pressure. Upon entry into the hippocampus (100-200 μm above the depth determined by the LFP measurement), the pressure was reduced to 0.15-0.30 psi. Cells were identified based on reproducible increases in electrode resistances, and then the whole-cell configuration was achieved.

Forty pyramidal neurons were classified as place cells based on their firing characteristics: firing rate in ten consecutive spatial bins (20 cm) exceeded 20% of the peak firing rate, and the mean in-field firing rate was more than three times the mean out-of-field firing rate. Of these, twenty-two were experimentally induced via somatic current injections (300 ms, 800 pA) applied during 6 consecutive laps at the same spatial location, as previously described ^68^. For MSOP and dextran experiments, the LFP electrode was filled with a solution containing MSOP (100 µM; Tocris, Minneapolis, MN) or Texas Red dextran 10 KDa (1 mM; Cat# D1828, ThermoFisher Scientific, Waltham, MA). In control experiments, the LFP electrode was only filled with 0.9% saline vehicle solution.

#### Histology

In some experiments, SR-101 (1 mM) was added to the LFP recording electrode, to estimate the diffusion distance of drugs pressure applied through the pipette. Mice were transcardially perfused within 5 min from the end of the electrophysiology experiments, using PBS (20 ml) followed by 4% PFA in PBS (20 ml). Extracted brains remained overnight in 4% PFA and were then rinsed twice and stored in PBS. Then, we prepared 100-μm-thick sagittal sections of PFA-fixed brains using a vibrating blade microtome (VT1000S; Leica Microsystems, Buffalo Grove, IL). The slices were incubated with streptavidin AF488 (1:750) and Triton X-100 0.2% overnight at 4°C, washed 3 times with PBS for 15 min, and mounted on glass slides using Fluoromount G DAPI mounting medium (Cat# 00-4959-52, ThermoFisher Scientific, Waltham, MA). All histological images were acquired on a Nikon Ti2E inverted confocal microscope equipped with 405/488/561/640 nm laser lines and Nikon Elements image acquisition software version 6.10.01.

#### Antibodies

Information about primary antibodies used for EM is reported below.

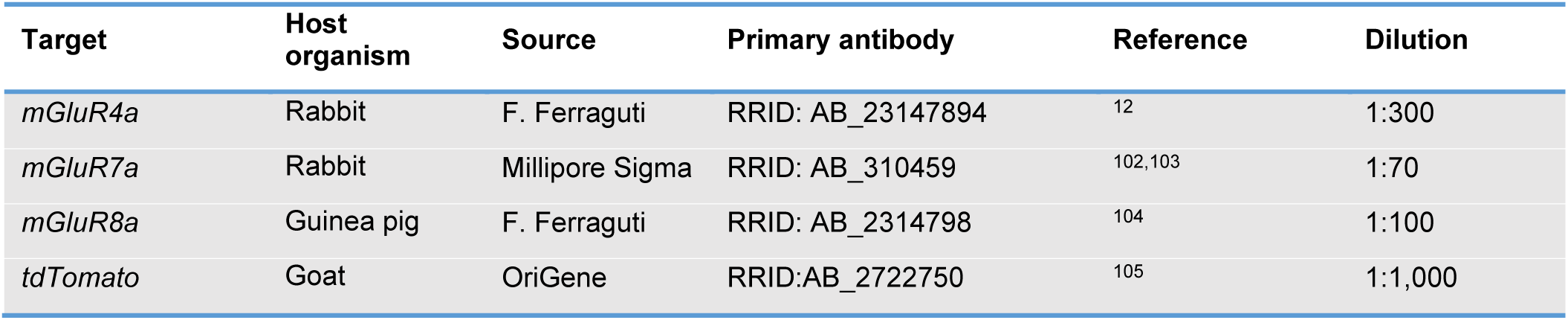

Information about secondary antibodies used for EM is reported below.

| Target | Host organism | Source | Secondary antibody | Reference | Dilution |
| --- | --- | --- | --- | --- | --- |
| *Rabbit IgG* | Donkey | Jackson Immunoresearch | RRID: AB_2340594 | - | 1:250 |
| *Guinea pig IgG* | Donkey | Jackson Immunoresearch | RRID: AB_2340452 | - | 1:250 |
| *Goat IgG* | Donkey | AURION | RRID:AB_2943420 | - | 1:40 |

#### Tissue collection and pre-embedding immunolabeling for EM analysis

The ultrastructural localization of mGluRIII receptors was analyzed using single immunoperoxidase EM in tissue samples from C57BL/6J (n=2) aged P40. Double immunoperoxidase EM experiments were performed in tissue collected from PV^Cre/+^:Ai9^Tg/Tg^ (n=2) and SST^Cre/+^:Ai9^Tg/Tg^ mice (n=2) aged P33-34. In both cases, mice were anesthetized with an intraperitoneal injection of chloral hydrate (4 mg/g, w/w; Cat# 07-894-1075; Patterson Veterinary, Loveland, CO) and perfused transcardially with a flush of 100 ml ice-cold, oxygenated ACSF (80 ml/min) containing (in mM): NaCl 119, KCl 2.5, CaCl_2_ 0.5, MgSO_4_ · H_2_O 1.3, MgCl_2_ 4, NaHCO_3_ 26.2, NaH_2_PO_4_ 1, glucose 22, pH 7.4 followed by 100 ml PFA4% in 0.1 M phosphate buffer (PB; pH 7.4; 80 ml/min). Brains were removed, post-fixed in the same fixative for 48 hrs at 4°C. They were washed and transferred to PBS added with an antiproteolytic cocktail (MG132 50 μM, pepstatin 1 μg/ml, leupeptin 20 μM, aprotinin 0.05 U/ml, and phenylmethylilphonyl fluoride 3.6 μM), and stored at 4°C for up to two weeks. Serial parasagittal sections (50 µm thick) were prepared using a vibrating blade microtome (PELCO easiSlicer, Ted Pella, Redding, CA) and collected in PB until processing ^106^.

For single immunoperoxidase pre-embedding studies, hippocampal sections were treated with H_2_O_2_ (1% in PB; 30 min) to remove endogenous peroxidase activity, rinsed in PB and pre-incubated in 10% Normal Donkey Serum (NDS, 1 hr). Sections were then incubated overnight at room temperature (RT) in a solution containing the primary antibody mixture. The following day, sections were rinsed 3 times in PB and incubated first in 10% NDS (15 min) and then in a solution containing the secondary biotinylated antibodies (1.5 hr at RT). Sections were subsequently rinsed in PB, incubated in avidin-biotin peroxidase complex (Vectastain Elite ABC-HRP kit, Cat# PK6100, Vector Laboratories, Newark, CA), washed several times in PB, and incubated in 3,3’diaminobenzidine tetrahydrochloride (DAB; 0.05% in 0.05 M Tris buffer, pH 7.6 with 0.03% H_2_O_2_). Method specificity was verified by substituting the primary antibodies with PB. After completion of the immunoperoxidase procedure, the sections were post-fixed in 1% osmium tetroxide in PB for 45 min and contrasted with 1% uranyl acetate in maleate buffer (pH 6.0; 1 hr) ^106^. The sections were then dehydrated in ethanol and propylene oxide, embedded in Epon/Spurr resin, flattened between Aclar sheets (Cat# 50426-25, Electron Microscopy Sciences, Hatfield, PA) and polymerized at 60°C for (48 hrs). Section chips including the pyramidal layer and *stratum radiatum* of hippocampal area CA1 were selected by light-microscopic inspection, glued to blank epoxy, and sectioned with an ultramicrotome (MTX, RMC Boeckeler, Tucson, AZ). The most superficial ultrathin sections (∼60 nm) were collected and mounted on copper grids, stained with Sato’s lead, and examined with a Philips CM100 transmission electron microscope coupled to a MegaView G2 TEM camera (Evident-Olympus, Center Valley, PA) controlled by software from Soft Imaging System GmbH, Muenster, Germany. To minimize the effects of procedural variables, material from each animal for each antigen was processed in parallel.

For double immunoperoxidase-immunogold labeling studies, following the completion of the immunoperoxidase procedure for mGluR4a, mGluR7a, or mGluR8a, sections were exposed to the pre-embedding immunogold silver enhancement procedure for the identification of of PV/SST-INs via detection of tdTomato in tissue collected from PV^Cre/+^:Ai9^Tg/Tg^ and SST^Cre/+^:Ai9^Tg/Tg^ mice^107^. Briefly, according to Yi *et al.* ^108^ and the manufacturer instructions (AURION, Wageningen, The Netherlands) after several rinses in PBS, sections were incubated with a tdTomato goat primary antibody (Cat# AB8181; SICGEN, Cantanhede, Portugal) dissolved in incubation buffer (PBS with 0.2% BSA-c^TM^, Cat# 900.022; AURION, Wageningen, The Netherlands) overnight at RT, washed in incubation buffer and then in colloidal gold ultrasmall conjugated anti-goat secondary antibody dissolved in incubation buffer (Cat# 800.333; AURION, Wageningen, The Netherlands) for 4 hrs at RT and overnight at 4°C. After several rinses in incubation buffer and PBS, the sections were immersed in 2.5% glutaraldehyde in PBS for 1 hr, rinsed several times in PBS and in Enhancement Conditioning Solution (ECS Cat# 500.055; AURION, Wageningen, The Netherlands), and then incubated with a silver enhancement reagent (R-Gent SE-EM Cat# 500.044; AURION, Wageningen, The Netherlands) for 60 min under dark conditions. After silver enhancement, the sections were osmicated in 0.5% osmium tetroxide in distillated water for 15 min, then rinsed in distilled water, PBS and last in maleate buffer. The sections were then contrasted with 1% uranyl acetate in maleate buffer (pH 6.0; 1 hr) and embedded in Epon/Spurr resin. Section chips including the pyramidal layer and *stratum radiatum* of hippocampal area CA1 were selected by light-microscopic inspection, glued to blank epoxy, and sectioned with an ultramicrotome. The most superficial ultrathin sections (∼60 nm) were collected and mounted on copper grids, stained with Sato’s lead, and examined with a Philips CM100 transmission electron microscope.

#### NEURON model of CA1-PCs

We built a compartmental model of CA1-PCs using the 3D morphology of a biocytin-filled CA1-PC generated previously by our lab (https://www.neuromorpho.org) (**Figure S7A**). All files pertaining to the NEURON simulations were uploaded to the ModelDB database (https://senselab.med.yale.edu/modeldb/, ModelDB acc.n. 000). The NEURON model was created to reproduce the results of the voltage-clamp experiments described in **Figure 5B-I** and was run on Microsoft Visual Studio Code as an IDE. In this model, the CA1-PC was voltage-clamped at 0 mV, and a leak potassium conductance (*g_pas_*) was used to reproduce space clamp errors associated with somatic voltage-clamp recordings ^64,109^ (**Figure S7B-D**). We used NRN-EZ (v1.1.6) ^110^ to randomly distribute 100 I-inputs along the soma and apical dendrites and set 𝑔_𝑝𝑎𝑠_ = 10^−5^(𝑒^𝑑⁄^_100_) (S/cm^2^), where *d*=distance between the soma and each compartment expressed in µm (**Figure S7B-D**).

Once the relationship between *g_pas_* and *d* was set, we performed separate simulations to constrain the synaptic weight of each inhibitory input. This was done empirically, working under the assumption that in many neocortical neurons voltage escape prevents recording the activity of synaptic inputs located >200 μm from the soma ^64,111,112^. This synaptic weight was set to 485 pS for both PV- and SST-inputs, to generate somatic mIPSCs with a mean amplitude of 12.8 pA, similar to the one recorded experimentally (12.7±1.0 pA, n=10; **Figure 3A-B; Figure S7E-F**). The local rise and exponential decay time of the I-inputs in the model were set to 1.5 and 20.0 ms, respectively, to ensure that the mIPSC kinetics at the soma matched those recorded experimentally (mIPSC rise model = 3.4 ms, t_50_ model = 17.8 ms; mIPSC rise exp = 3.6±0.1 ms, t_50_ exp = 17.8±0.9 ms n=10; **Figure 3A-B**). We constrained the release probability of PV- and SST-INs synapsing onto CA1-PCs based on previous work indicating that *Pr*_PV_=0.8 at [𝐶𝑎^2+^]_𝑜_=2 Mm ^66^ (**Figure S7G**). Given that there is a 4^th^ power relationship between *Pr* and [𝐶𝑎^2+^]_𝑜_ ^19,65^, we estimated *Pr*_PV_=0.1 at [𝐶𝑎^2+^]_𝑜_=1.2 mM, which we used in our recording conditions (**Figure S7G**). Based on our optogenetics experiments, the number of active PV release sites required to evoke an oIPSC comparable to the oIPSC evoked experimentally is 4. Therefore, we estimated a total of 4/0.1=40 PV release sites targeted CA1-PC in our model. EM data indicate that there are 8.3 times more SST- than PV-inputs onto CA1-PCs ^67^. Therefore, the total number of SST release sites was estimated as 40⋅8.3=332. The number of active SST sites required to generate an oIPSC with the same amplitude as the oIPSC recorded experimentally was 18, leading to *Pr*_SST_=18/332=0.05 (**Figure S7G**). Each simulation was repeated 100 times, each time randomizing the location of the active inputs. Therefore, the results represent the average of 100 simulations.

#### EM analysis

Immuno-positive profiles were studied from ultrathin sections at the surface of the embedded blocks. Quantitative data on profiles derived from the analysis of hippocampal neuropil (10–15 ultrathin sections/mouse; 2 mice for each antigen). Fields of view with mGluR4a, mGluR7a, and mGluR8a positive profiles were randomly selected, and captured at 19,000-34,000× magnification. Acquisition of microscopical fields and analysis were performed under blinded conditions. For the analysis of the distribution of mGluRIII positive profiles, subcellular compartments (i.e., dendrites, axons, axon terminals, and astrocytic processes) were identified according to well-established criteria ^113^. In line with previous quantitative EM of synapses^114^, in a synapse the pre-synaptic terminal was characterized by vesicles adjacent to the pre-synaptic density, the synaptic cleft displayed electrodense material, the pre- and post-synaptic membranes defining the pre- and post-synaptic specializations were characterized by electron densities, and finally the prominent and/or thin post-synaptic density in addition to clear and round or pleiomorphic vesicles permitted to distinguish asymmetric and symmetric synapses ^106,114^. According to different post-synaptic targets of interneurons, immuno-positive axon terminals forming symmetric synaptic contacts were sampled at axo-somatic, proximal, distal, or spinous axo-dendritic synapses (dendrites were considered distal if their diameter was ≤1 μm, proximal if it was >1 μm ^115–119^. Astrocytic processes were identified by their typical irregular outlines, the paucity of cytoplasmic components (except for ribosomes, glycogen granules, and some bundles of fibrils^113^. Pre- embedded double labeled material, EM inspection of ultrathin sections (10-12 per mouse/antigen) allowed identifying axon terminals of PV- and SST-INs based on the presence of silver amplified signal within terminals synapsing with post-synaptic domains and to recognize electron-dense immunoreacted products relative to mGluRIII isoforms in the close vicinity of the active zone.

#### In vitro electrophysiology analysis

Electrophysiological recordings were analyzed within the Igor Pro environment using custom made software (A. S.). EM data were analyzed within the GraphPad Prism environment.

#### In vivo electrophysiology and behavior analysis

We analyzed the *in vivo* electrophysiology and behavioral data using custom code written in MATLAB 2023b (Mathworks, Natick, MA). The analysis was performed on periods of time when the mouse was running at a speed >4 cm/s. APs were detected using the first derivative of the raw V_m_, as the time points when its depolarization rate crossed a threshold of 40 mV/ms. To analyze the slow V_m_, we removed APs from the raw traces, replaced them using a linear interpolation of the raw V_m_, down-sampled the resulting V_m_ to 200 Hz, and then low-pass filtered them at 2 Hz using a finite impulse response (FIR) filter with a 2 s Hamming window. To ensure that V_m_ and slow V_m_ had the same number of data points, we resampled the slow V_m_ at 20 kHz. The baseline value of the slow V_m_ was defined as the mode of the slow V_m_ calculated over the entire recording. This value was subtracted from the slow V_m_ to calculate the absolute amplitude of the V_m_ ramp depolarization for each bin. In-field locations extended for a total of 20 cm (i.e., ten bins before and ten bins after the peak V_m_). To calculate the intracellular theta amplitude, we applied a Hilbert transform to the subthreshold V_m_ (i.e., the AP-removed but unfiltered V_m_), after it was down- sampled to 200 Hz, band-pass filtered at 4-7 Hz (2 s Hamming window) and resampled at 20 kHz. The AP frequency, slow V_m_, baseline V_m_-corrected V_m_, and intracellular theta amplitude were calculated over 2 cm bins (10 bins located around the peak and 10 bins located at the end of the out-of-field locations) and averaged across 20- 40 laps. Plateau potentials were defined as times of prolonged depolarization (>-35 mV for >10 ms), identified using a boxcar running average of the subthreshold V_m_ (401 points, ∼20 ms).

### Quantification and statistical analysis

Data are presented as mean ± SEM unless otherwise stated. Normality was tested using SPSS or GraphPad Prism 8.2.1. Statistical significance was determined by Student’s paired or unpaired t test, ANOVA, General Estimated Equations (**Figure 2U-V**; **Figure 6C, F, I, L; Figure S9B, E, H, K**) or Fisher exact test (EM), as appropriate (IgorPro 6.37, SPSS, GraphPad Prism 8.2.1). Differences were considered significant at p<0.05 (*p<0.05; **p<0.01; ***p<0.001).

