## Supplementary figures and images for "An unsuspected physiological role for mGluRIII glutamate receptors in hippocampal area CA1"

### Fig S1

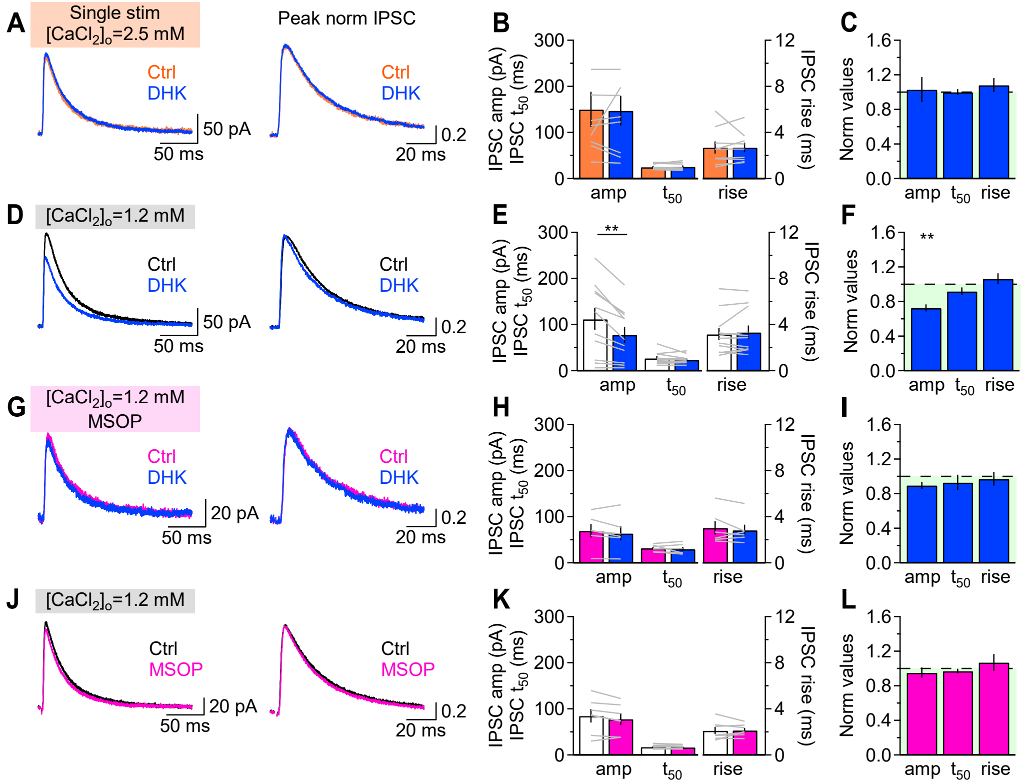

### Fig S2

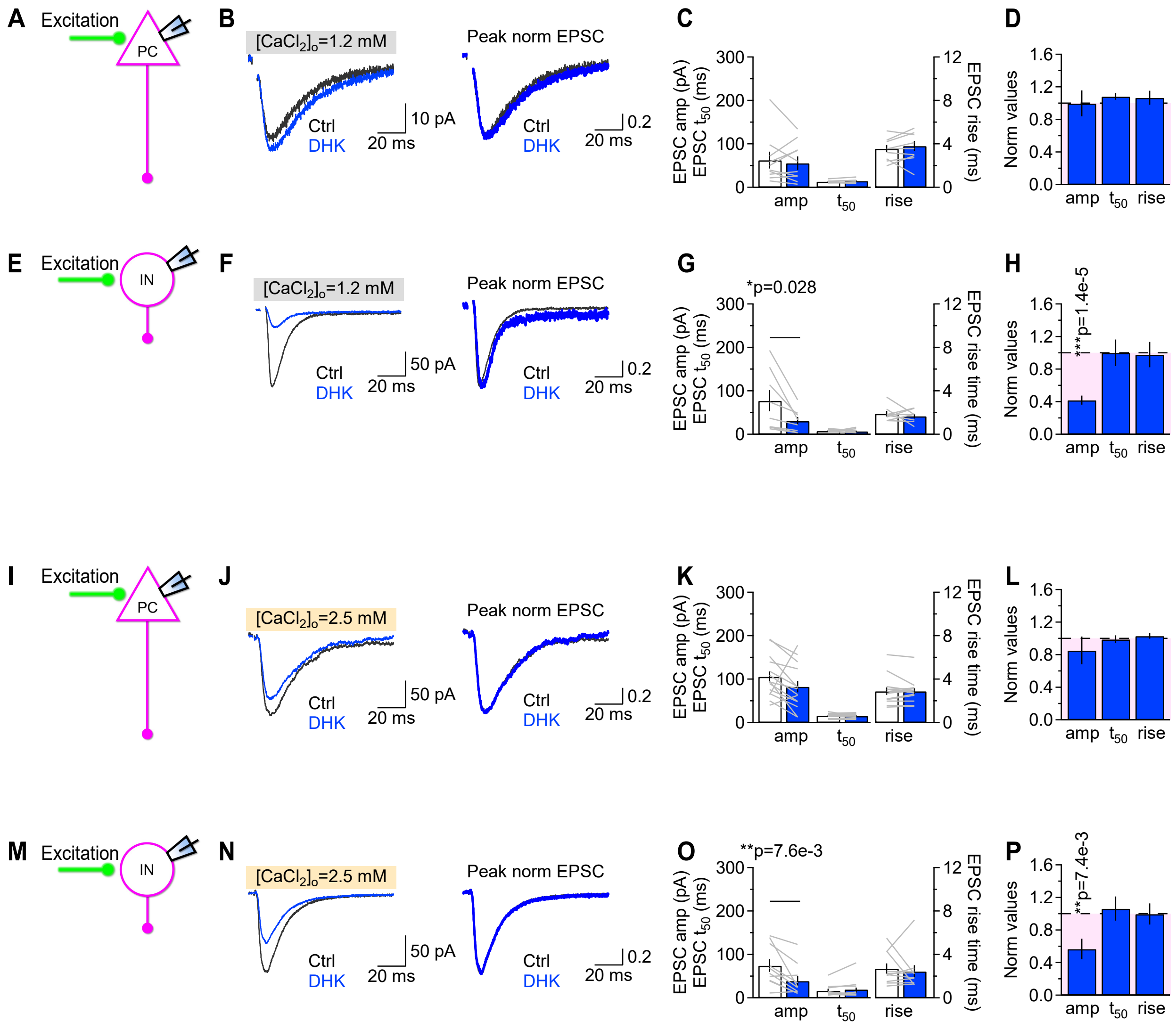

### Fig S3

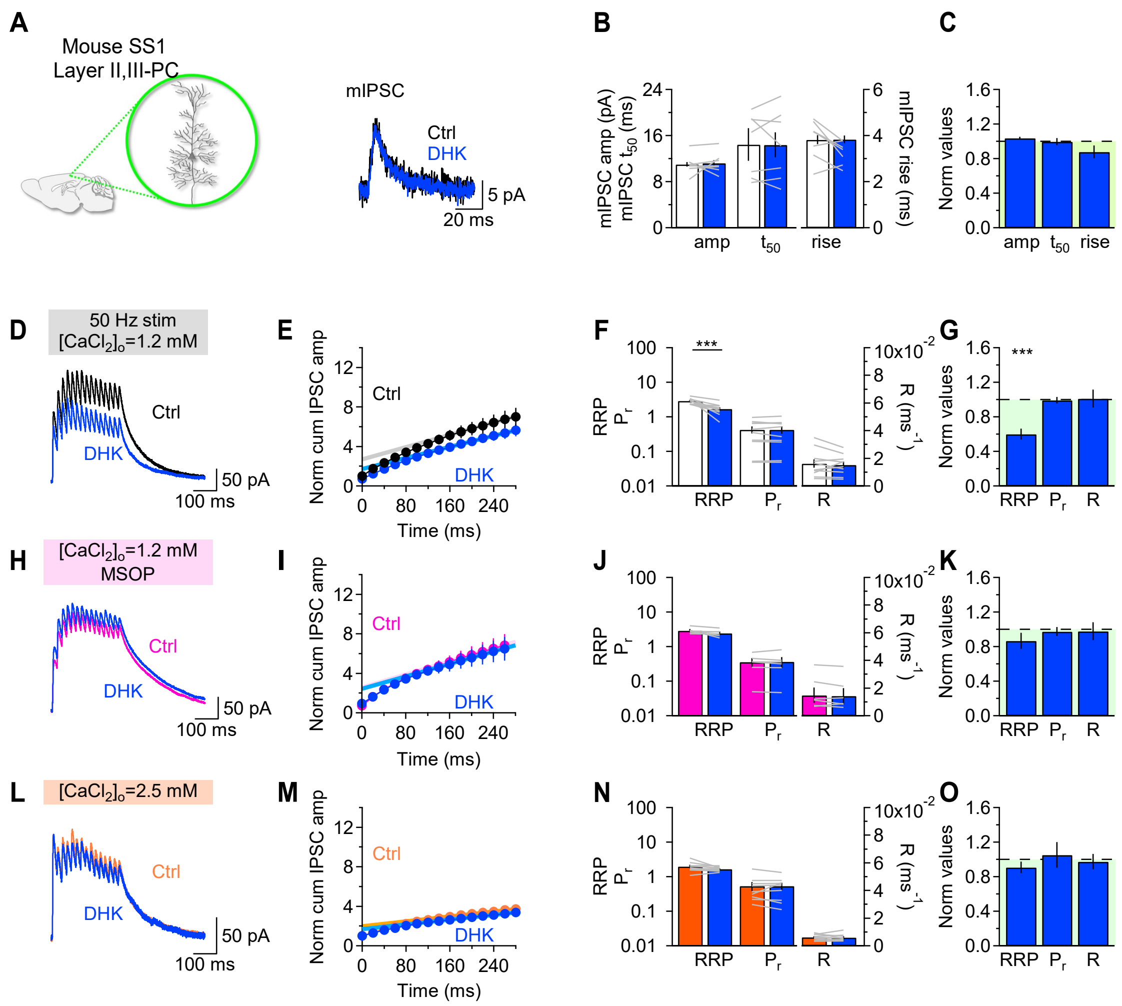

### Fig S4

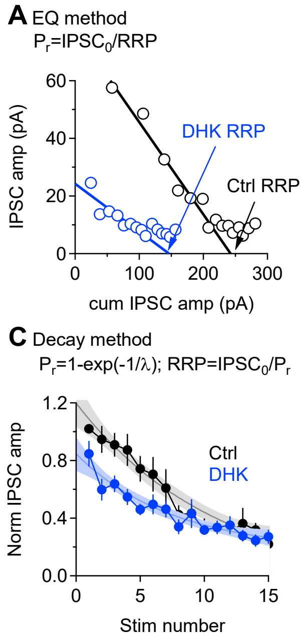

### Fig S5

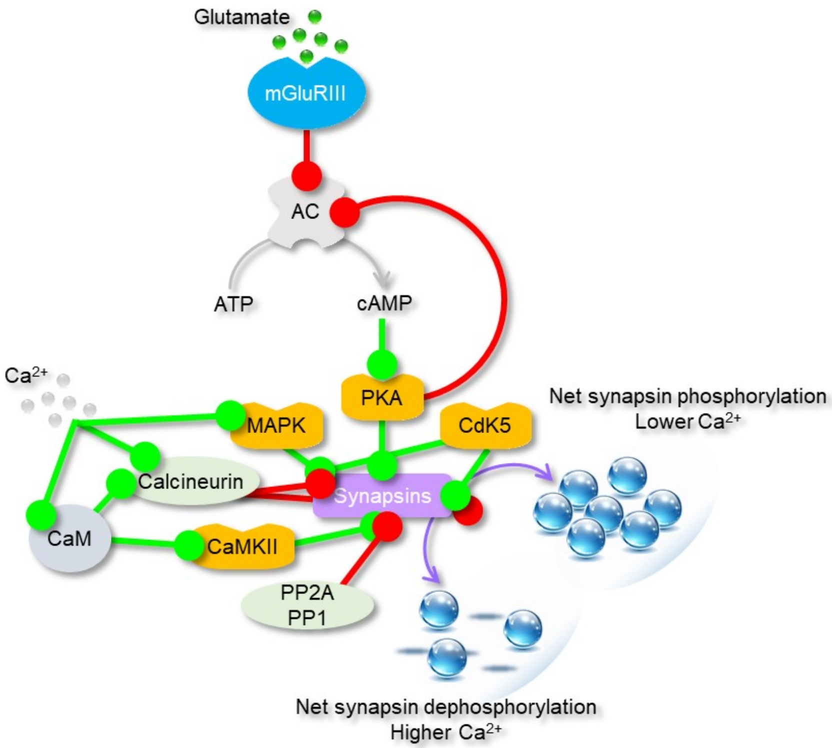

### Fig S6

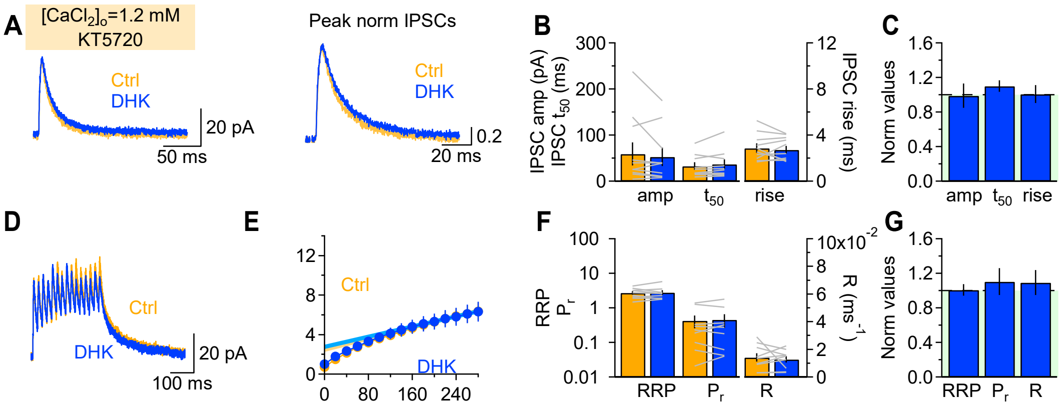

### Fig S7

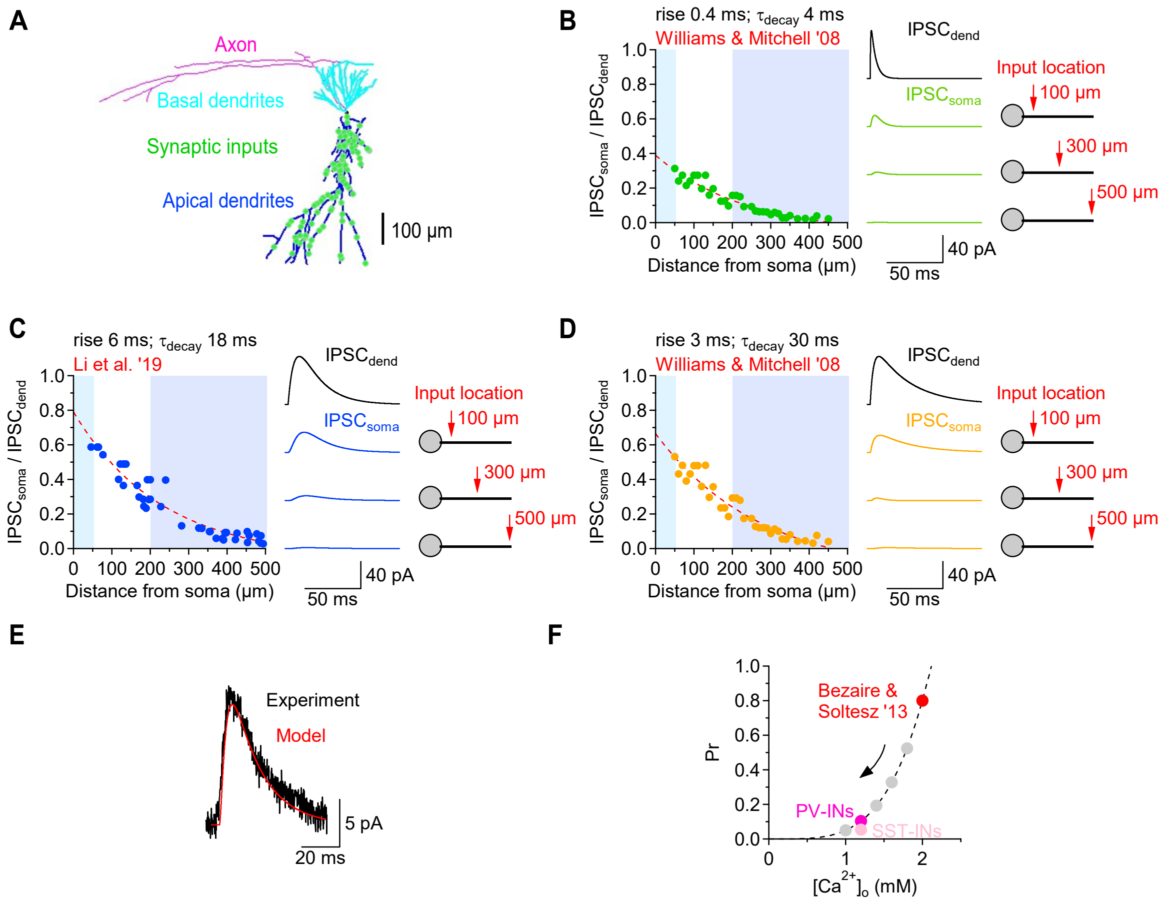

### Fig S8

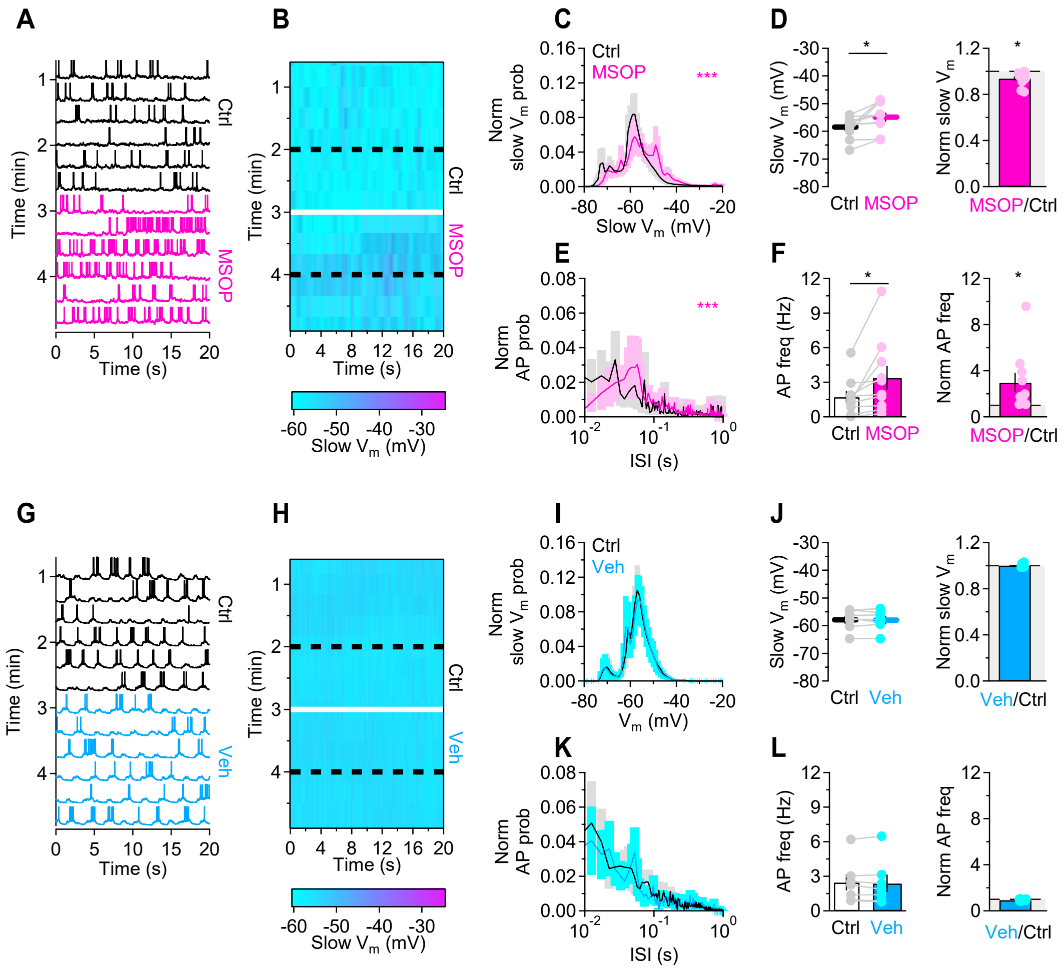

### Fig S9

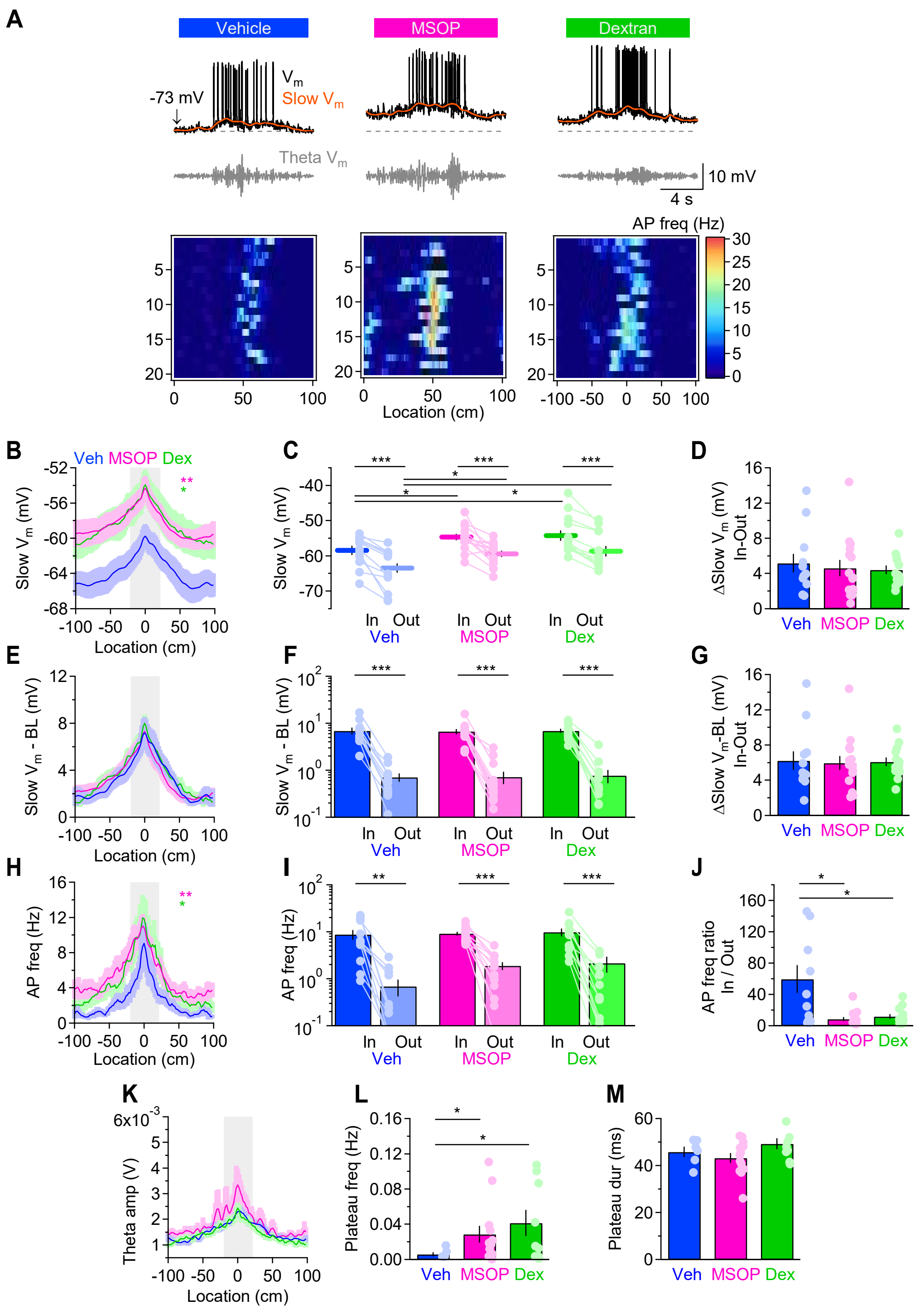
