## Supplementary material for "An unsuspected physiological role for mGluRIII glutamate receptors in hippocampal area CA1": Supp legends

### Supplementary Figure Legends

**Figure S1. mGluRIII activation reduces inhibition onto SS1-PCs. (A)** Average IPSCs recorded with $[Ca]_o^{2+}$=2.5 mM in SS1-PCs, before (*orange*) and after DHK (100 µM; *blue*). The traces on the right are normalized by the peak IPSC amplitude. **(B)** Summary of the effect of DHK on the IPSC amplitude and kinetics. **(C)** Summary of the relative change in IPSC amplitude and kinetics induced by DHK. **(D-I)** As in A-C, DHK was applied in the presence of $[Ca^{2+}]_o$=1.2 mM (D-F), and in the continued presence of MSOP (100 µM; G-I). **(J-L)** As in A-C, in baseline conditions and in the presence of MSOP.

**Figure S2. In physiological calcium, mGluRIII activation reduces inhibition onto CA1-INs, not CA1-PCs. (A)** Schematics of synaptic connections tested in these experiments. **(B)** Average EPSCs recorded with $[Ca^{2+}]_o$=1.2 mM in CA1-PCs, before (*black*) and after DHK (100 µM; *blue*). The traces on the right are normalized by the peak EPSC amplitude. **(C)** Summary of the effect of DHK on the EPSC amplitude and kinetics. **(D)** Summary of the relative change in EPSC amplitude and kinetics induced by DHK. **(E-H)** as in A-D, for EPSCs recorded from CA1-INs in control conditions and in the presence of DHK. **(J-P)** as in A-H, for EPSCs recorded from CA1-PCs (I-L) and INs (M-P) in $[Ca^{2+}]_o$=2.5 mM.

**Figure S3. Different techniques confirm that glutamate spillover reduces the RRP at I-synapses onto CA1-PCs. (A)** Representative recording from which we estimated *Pr* and *RRP* using the EQ method [27]. At the beginning of the train stimulation, there is a virtually linear fall of the IPSC amplitude with respect to the cumulative IPSC. This can be expected if a constant fraction of the RRP was being released, and vesicle refiling was negligible. **(B)** Summary showing that DHK reduced the *RRP* without altering *Pr*. **(C)** Estimates of *Pr* and *RRP* using the decay method [34]. According to this approach, the amplitude of the n$^{th}$ IPSC in a train is $IPSC_n = IPSC_0(1 - P_r)^n + K$, where $K$ is a constant. An exponential fit to the IPSC amplitude as a function of the stimulus number can be used to estimate *Pr*, and $RRP = IPSC_o/P_r$. **(D)** Summary graph showing that the decay method confirms that DHK reduces the *RRP* without altering *Pr*.

**Figure S4. mGluRIII activation reduces the size of the *RRP* at I-synapses onto SS1-PCs. (A)** *Left,* Schematic representation of SS1-PCs in sagittal brain slices. *Right,* Average mIPSCs recorded in a SS1-PC in control conditions (*black*) and in DHK (100 µM; *blue*). **(B)** Mean values and in-cell comparison of the effect of DHK on the mIPSC amplitude and kinetics. **(C)** Summary of the effect of DHK on the mIPSC amplitude and kinetics, normalized by their values in control conditions. **(D)** Representative IPSC train recorded in response to cortical stimulation with 15 pulses at 50 Hz (280 ms), before (*black*) and after DHK (*blue*), in $[Ca^{2+}]_o$=1.2 mM. **(E)** Cumulative IPSC amplitude normalized by the amplitude of the first IPSC in the train, plotted as a function of the stimulation timing, in control conditions (*black*) and in DHK (*blue*). Thich lines represent the linear fit of the last 5 IPSCs in each condition. The y-intercept of these lines was used to estimate the *RRP*. **(F)** Mean and in-cell comparison of the effect of DHK on *RRP*, *Pr* and *R*. **(G)** Summary of the effect of DHK on *RRP*, *Pr* and *R*, normalized by their values in control conditions. **(H-K)** As in D-G, for recordings obtained in the continued presence of the mGluRIII antagonist MSOP (100 µM). **(L-O)** As in D-G, for recordings obtained in $[Ca^{2+}]_o$=2.5 mM.

**Figure S5. Signaling cascades involved in the regulation of the RRP size.** The scheme summarizes signaling cascades coupled to mGluRIII and triggered by $Ca^{2+}$, which alter the phosphorylation state of synapsins, with consequences for the regulation of the size of the RRP.

**Figure S6. PKA inhibition prevents the spillover-induced reduction of the IPSC amplitude and *RRP* size at I-synapses onto SS1-PCs. (A)** Average IPSCs recorded with $[Ca^{2+}]_o$=1.2 mM in SS1-PCs, before (*orange*) and after DHK (100 µM; *blue*), in the continued presence of the PKA inhibitor KT5720 (1 µM). The traces on the right are normalized by the peak IPSC amplitude. **(B)** Summary of the effect of DHK on the IPSC amplitude and kinetics in KT5720. **(C)** Summary of the relative change in IPSC amplitude and kinetics induced by DHK applied in the continued presence of KT5720. **(D)** Representative IPSC train recorded in response to cortical stimulation with 15 pulses at 50 Hz (280 ms), before (orange) and after DHK (blue), in the continued presence of KT5720 and in $[Ca^{2+}]_o$=1.2 mM. **(E)** Cumulative IPSC amplitude normalized by the amplitude of the first IPSC in the train, plotted as a function of the stimulation timing, in control conditions (*orange*) and in DHK (*blue*). Thich lines represent the linear fit of the last 5 IPSCs in each condition. The y-intercept of these lines was used to estimate the *RRP*. **(F)** Mean and in-cell comparison of the effect of DHK on *RRP*, *Pr* and *R*, when applied in the presence of KT5720. **(G)** Summary of the effect of DHK on *RRP*, *Pr* and *R*, when applied in the continued presence of KT5720. The data were normalized by their values in control conditions.

**Figure S7. Parameter optimization for voltage-clamp model of CA1-PCs. (A)** Morphology of the CA1-PC used to run the NEURON compartmental model, with 100 inputs distributed randomly throughout the soma and apical dendrites (*green dots*). **(B-D)** Quantification of the distance-dependent loss of current at the soma, for IPSCs generates at different dendritic locations. Introducing a leak conductance $g_{pas} = 10^{-5}(e^{d/100})$ allows reproducing the somatic voltage clamp loss reported in the literature using dendritic recordings [64,109]. The traces represent the inhibitory currents injected in dendrites located 100, 300 and 500 µm away from the soma (*black*) and their corresponding somatic recordings (*green, blue, orange).* **(E)** Comparison of somatic mIPSCs recorded experimentally and generated through the model. **(F)** Power relationship describing the $Ca^{2+}$-sensitivity of the release probability[19]. The red dot represents the value of *Pr* of PV-to-CA1-PC inputs estimated in previous work with $[Ca^{2+}]_o$=2.0 mM [66]. The relationship was used to estimate *Pr* for PV-to-CA1-PC inputs in our recording conditions, where $[Ca^{2+}]_o$=1.2 mM (*magenta*). *Pr* for SST-inputs was calculated by dividing the number of estimated SST inputs by the number of active inputs that were recruited to mimic the oIPSCs recorded experimentally (*pink*).

**Figure S8. mGluRIII activation leads to membrane potential hyperpolarization of CA1-PCs in anesthetized mice. (A)** Raw membrane potential ($V_m$) recordings obtained 3 min before and after pressure application of MSOP (100 µM; *magenta*). **(B)** Representation of low-pass filtered, slow $V_m$. The white line represents the time of the MSOP application. **(C)** Probability distribution of low-pass filtered slow $V_m$ values recorded 1 min before and 1 min after MSOP (n=10 CA1-PCs, N=8 mice). The data are normalized by the area under the curve of the probability distribution in control conditions (*black*) and in MSOP (*magenta*). **(D)** *Left,* Average low-pass filtered slow $V_m$ recorded in control conditions and in MSOP. *Right,* In-cell comparison of slow $V_m$ before and after MSOP application. **(E)** Probability distribution of inter-spike interval (ISI) measured 1 min before and 1 min after MSOP (n=10 CA1-PCs, N=8 mice). The data are normalized by the area under the curve of the probability distribution in control conditions and in MSOP. **(F)** *Left,* Average AP frequency recorded in control conditions and in MSOP. *Right,* In-cell comparison of AP frequency following MSOP application. **(G-L)** Same as in A-F for control experiments in which a vehicle solution was applied through the LFP recording pipette (n=7 CA1-PCs; N=6 mice).

**Figure S9. Reducing spillover reduces the excitability of CA1-PCs in awake mice. (A)** Example of raw $V_m$ (*black*), low-pass filtered $V_m$ (slow $V_m$; *orange*), and theta-filtered intracellular $V_m$ (*gray*) for CA1-PC place cells recorded in vehicle, MSOP (100 µM) and dextran 10 KDa (1 mM). The heatmaps represent the spatial firing rate of these cells, calculated over 20 consecutive laps. The dashed line marks -73 mV. **(B)** Slow $V_m$ ramp of CA1 place cells. The gray shaded area represents the location of the in-field region. **(C-D)** Summary of the in-field, out-of-field, and the difference in the in-field vs. out-of-field slow $V_m$ values in vehicle (*blue*), MSOP (*magenta*) and dextran (*green*). MSOP and dextran increase the in-field and out-of-field slow $V_m$ by ~5mV. **(E)** Baseline (BL) $V_m$-subtracted slow $V_m$ showing no change in the magnitude of the in-field and out-of-field value between vehicle and MSOP. **(F-G)** Summary of the baseline subtracted in-field, out-of-field, and the difference in the in-field vs. out-of-field baseline-subtracted slow $V_m$ values. MSOP and dextran do not change the amplitude of the baseline $V_m$-subtracted slow $V_m$. **(H)** Spatial dependence of AP frequency. **(I-J)** Summary of the in-field, out-of-field, and in-to-out-of-field AP frequency ratio. MSOP and dextran decrease the in-field to out-of-field AP frequency ratio. **(K)** Spatial dependence of the amplitude of the intracellular theta activity. **(L-M)** Summary graphs of the plateau frequency and duration, respectively. MSOP and dextran increase the plateau frequency without altering the plateau duration.

### Supplementary Table Legends

**Supplementary Table 1. Distribution of mGluR4a, mGluR7a, and mGluR8a immunoreactivity in hippocampal area CA1.** Data were obtained from pre-embedded CA1-PC and *stratum radiatum* layers. Distribution among immunoreactive (IR) profiles is expressed in percentage. Abbreviations: number of IR profiles sampled (*n*); axon terminals with pleiomorphic vesicles forming symmetric synaptic contact onto CA1-PC somata (*AxT→Soma Sym*); axon terminals with pleiomorphic vesicles forming symmetric synaptic contact onto proximal dendrites of principal cells (*AxT→Prox Sym*); axon terminals with pleiomorphic vesicles forming symmetric synaptic contact onto distal dendrites or spines (*AxT→Dist Sym*); axon terminals containing exclusively clear round vesicles and forming asymmetric synaptic contacts (*AxT Asym*); axon terminals not making a recognizable synaptic contact (*AxT*); axons (*Ax*); proximal dendrites (*Prox*); distal dendrites (*Dist*); dendritic profiles (*Den*); astrocytic processes (*Astro*); unidentified profiles (*N/A*).

| Target | IR profiles | AxT→Soma sym | AxT→Prox sym | AxT→Dist sym | AxT asym | AxT | Ax | Den | Astro | N/A |
|---|---|---|---|---|---|---|---|---|---|---|
| *mGluR4a* | n=905 | 6% n=52 | 4% n=40 | 5% n=49 | 11% n=97 | 9% n=83 | 12% n=106 | 50% n=449 | 2% n=17 | 1% n=12 |
| *mGluR7a* | n=1056 | 5% n=48 | 4% n=42 | 6% n=64 | 8% n=89 | 10% n=107 | 11% n=119 | 48% n=503 | 6% n=64 | 2% n=20 |
| *mGluR8a* | n=472 | 6% n=27 | 3% n=14 | 5% n=23 | 10% n=47 | 8% n=36 | 7% n=31 | 57% n=270 | 4% n=17 | 2% n=7 |
